# Axonal transport of autophagosomes is regulated by dynein activators JIP3/JIP4 and ARF/RAB GTPases

**DOI:** 10.1101/2023.01.28.526044

**Authors:** Sydney E. Cason, Erika L.F. Holzbaur

## Abstract

Neuronal autophagosomes, “self-eating” degradative organelles, form at presynaptic sites in the distal axon and are transported to the soma to recycle their cargo. During transit, autophagic vacuoles (AVs) mature through fusion with lysosomes to acquire the enzymes necessary to breakdown their cargo. AV transport is driven primarily by the microtubule motor cytoplasmic dynein in concert with dynactin and a series of activating adaptors that change depending on organelle maturation state. The transport of mature AVs is regulated by the scaffolding proteins JIP3 and JIP4, both of which activate dynein motility in vitro. AV transport is also regulated by ARF6 in a GTP-dependent fashion. While GTP-bound ARF6 promotes the formation of the JIP3/4-dynein-dynactin complex, RAB10 competes with the activity of this complex by increasing kinesin recruitment to axonal AVs and lysosomes. These interactions highlight the complex coordination of motors regulating organelle transport in neurons.

**Summary:** Mature autophagosomes in the axon are transported by the microtubule motor dynein, activated by JNK-interacting proteins 3 and 4 (JIP3/4). This motility is regulated by the small GTPases ARF6 and RAB10. The tight regulation of autolysosomal transport is essential for intracellular recycling to maintain neuronal homeostasis.

## Introduction

Maintaining neuronal homeostasis across the lifespan requires the continuous turnover of dysfunctional or aged proteins and organelles (Eskelinen, 2019; Kulkarni et al., 2018). Autophagy is a process by which these components can be broken down and recycled (Stavoe and Holzbaur, 2019). Autophagic vacuoles (AVs), the “self-eating” organelle, engulf cargo proteins or organelles in a double-membrane and then fuse with late endosomes and lysosomes (collectively, endolysosomes), which provide the degradative enzymes necessary to breakdown the cargo (Yim and Mizushima, 2020). In neurons, AVs form preferentially at presynaptic sites and at the distal tip of the axon and must be actively transported to the soma, where the majority of protein and organelle biogenesis occurs (Maday et al., 2012; Maday and Holzbaur, 2014; Stavoe et al., 2016; Koltun et al., 2020; Farfel-Becker et al., 2019). The transport of AVs along the axon is primarily driven by the microtubule motor cytoplasmic dynein I, in coordination with its obligate partner complex dynactin (Kimura et al., 2008; Katsumata et al., 2010; Maday et al., 2012). The dynein-dynactin complex needs to be recruited to and activated locally on the AV by adaptor proteins (Fu et al., 2014; Cheng et al., 2015; Khobrekar et al., 2020; Cason et al., 2021). The opposing motor kinesin-1 also localizes to axonal AVs where it may compete with dynein (Wong and Holzbaur, 2014; Maday et al., 2012). Kinesin inactivation on AVs is essential for autophagic transport and flux, and its dysregulation can be observed in the context of neurodegenerative disease (Fu et al., 2014; Boecker et al., 2021; Dou et al., 2022).

The simultaneous activation of dynein-dynactin and inactivation of kinesin must therefore be coordinated locally at the AV membrane. This regulation is further complicated by autophagosomal maturation, during which the AV membrane and associated proteins are altered via fusion with endolysosomes (Cason et al., 2021). We previously found that different motor regulatory proteins drive the retrograde transit of AVs along the axon, dependent upon the sub-axonal location and maturation state of the AV (Cason et al., 2021). Specifically, JNK-interacting protein (JIP) 1 regulates the initial transit of nascent AVs in the distal axon by inactivating kinesin-1 (Fu et al., 2014). Huntingtin-associated protein 1 (HAP1) activates dynein on partially mature AVs in the mid-axon (Wong and Holzbaur, 2014; Cason et al., 2021). Finally, the motility of the most mature population of axonal AVs is regulated by the motor-interacting protein JIP3 (Cason et al., 2021).

Mutations in the JIP3 gene (*MAPK8IP3;* homolog of UNC-16 and Sunday Driver/Syd) result in a rare neurodevelopmental disorder, and JIP3 expression is relatively limited to the brain; in contrast, the related protein JIP4 (*SPAG9*) is expressed ubiquitously (Ito et al., 1999; Jagadish et al., 2005; Kelkar et al., 2000; Platzer et al., 2019). Recent work has shown that JIP3 and JIP4 contain a structurally conserved motif in their N-termini that mediates binding to dynein light intermediate chain (DLIC), a feature common among dynein activating adaptors (Celestino et al., 2022). Further, JIP3/4 can bind to the dynactin subunit p150^Glued^ (Fig. 1 A) and truncated JIP3 can activate dynein motility in a purified system (Montagnac et al., 2009; Rao et al., 2022). However, JIP3 and JIP4 can also interact with the kinesin-1 complex via interactions (Fig. 1 A) with kinesin heavy chain (KIF5) and kinesin light chain (KLC) (Arimoto et al., 2011; Cavalli et al., 2005; Celestino et al., 2022; Cockburn et al., 2018; Montagnac et al., 2009; Sun et al., 2011; Tuvshintugs et al., 2014; Vilela et al., 2019). Further, JIP3 has been reported to induce kinesin activity *in vitro* (Sun et al., 2011; Watt et al., 2015).

**Figure 1.**
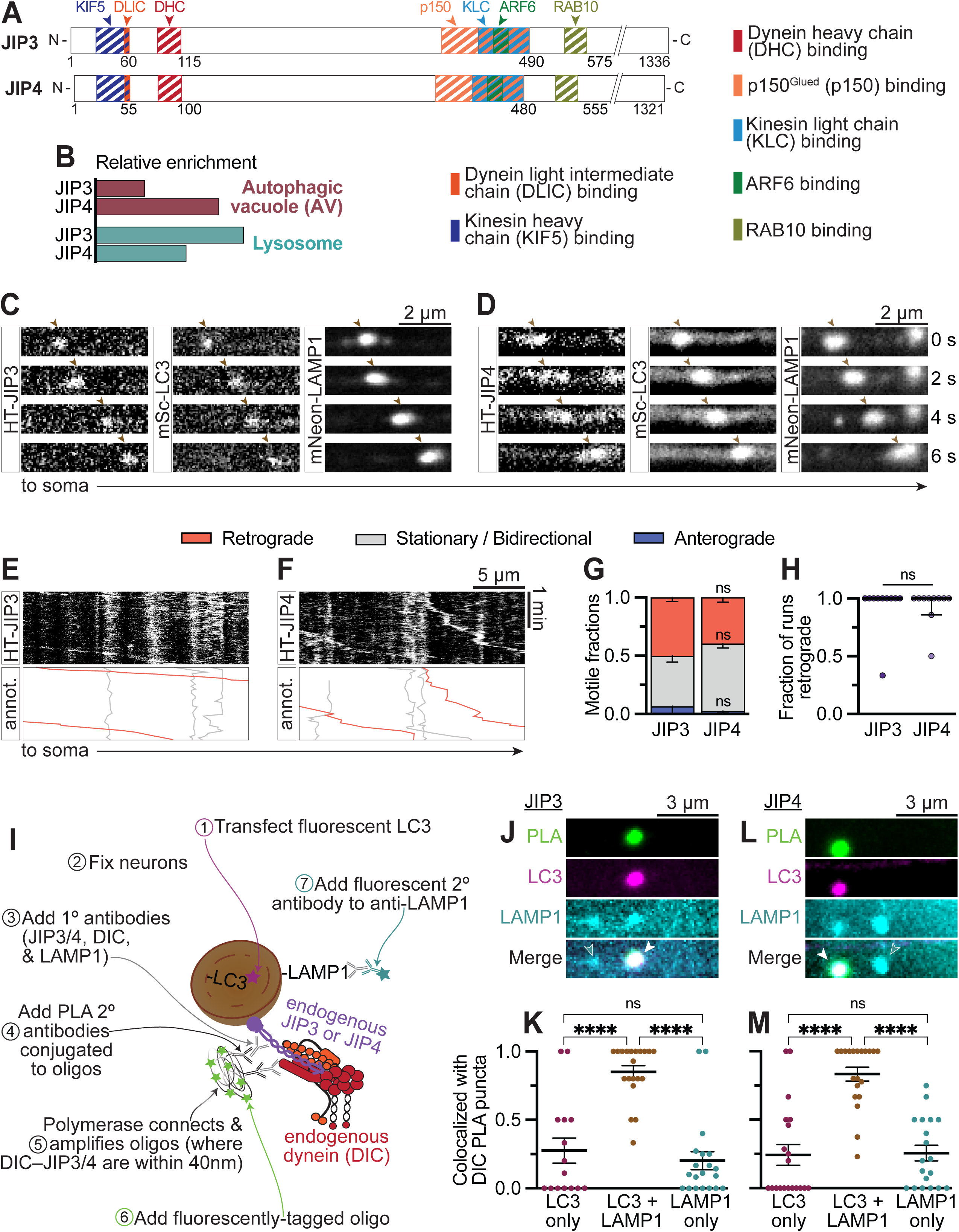
JIP3/4 comigrate with and interact with dynein on autolysosomes. **(A)** N-terminal region of JIP3 and JIP4, scaled to the primary sequence. Arrows: green, JIP-DIC PLA only; magenta, LC3 only; cyan, LAMP1 only; ochre, LC3 + LAMP1; white, LC3 + LAMP1 + JIP-DIC PLA. **(B)** Relative enrichment (normalized number of peptides, see Methods for details) for JIP3 (Mapk8ip3) and JIP4 (Spag9) in the proteomics performed by Goldsmith et al., (2022) and Dumrongprechachan et al., (2022). **(C-D)** Time series demonstrating JIP3 and JIP4 comigration with LC3 (AV marker) and LAMP1 (lysosome marker). **(E-F)** Example kymographs from the proximal axons of neurons transfected with JIP3 or JIP4. Kymographs depict distance on the x-axis and time on the y-axis. Annotated kymographs (annot.) mirror the above kymographs with the JIP3/4+ puncta paths pseudo-colored for visualization. **(G)** Quantification of JIP3/4+ puncta moving retrograde (≥10µm towards the soma), anterograde (≥10µm towards the axon tip), or exhibiting bidirectional/stationary motility (moving <10µm). n = 10 neurons; two-way ANOVA with Sidak’s multiple comparisons test (anterograde, P = 0.8976; stationary/bidirectional, P = 0.0767; retrograde, P = 0.2792). **(H)** Fraction of motile events (≥ 10 µm either direction) moving retrograde. n = 10 neurons; unpaired t test (P = 0.9776). **(I)** Schematic illustrating the proximity ligation assay (PLA). **(J-M)** Example micrographs and quantifications showing colocalization between LC3, LAMP1, and JIP3-DIC (J-K) or JIP4-DIC (L-M) puncta. n = 20 neurons; one-way ANOVA with Tukey’s multiple comparisons test; JIP3 (LC3 v. LC3 + LAMP1, P < 0.0001; LC3 v. LAMP1, P = 0.7236; LAMP1 v. LC3 + LAMP1, P < 0.0001); JIP4 (LC3 v. LC3 + LAMP1, P < 0.0001; LC3 v. LAMP1, P = 0.9879; LAMP1 v. LC3 + LAMP1, P < 0.0001).

We therefore asked how JIP3 and JIP4 are regulated to drive the highly processive retrograde transit of AVs along the axon. JIP3 and JIP4 both associate with mature AVs as well as lysosomes, and either full-length protein can activate dynein motility in an *in vitro* assay. We used proteomic databases to identify potential regulators of JIP3/4-dependent motility, and found that the small GTPase ARF6 is enriched in AVs isolated from brain (Goldsmith et al., 2022). In live neurons, we demonstrate that the GTPase state of ARF6 is important for the regulation of both AV and lysosomal motility along axons. Another JIP3/4-interacting GTPase, RAB10, enriched in a lysosomal fraction from brain (Dumrongprechachan et al., 2022), also affected AV and lysosomal motility along axons, but in a distinct fashion. We therefore propose that the recruitment and activation of JIP3/4 at organellar membranes is differentially regulated by discrete small GTPases to generate unique motile behaviors. JIP3/4 represent a growing group of motor-activating proteins that can bind both dynein and kinesin motors on organelle cargos (Arimoto et al., 2011; Bielska et al., 2014; Canty et al., 2021; Cason et al., 2021; Celestino et al., 2022; Colin et al., 2008; Fenton et al., 2021; Fu and Holzbaur, 2013; Kendrick et al., 2019; López-Doménech et al., 2018; Twelvetrees et al., 2019; Vilela et al., 2019; Zhao et al., 2021), and must be tightly regulated by additional binding partners to induce unidirectional transport in the cell.

## Results

### JIP3 and JIP4 interact with dynein on autolysosomes

Previous studies have implicated both JIP3 and JIP4 in the transport of a number of organelles, especially degradative vesicles such as AVs, endosomes, and lysosomes (Abe et al., 2009; Boecker et al., 2021; Brown et al., 2009; Cason et al., 2021; Choudhary et al., 2017; Drerup and Nechiporuk, 2013; Hill et al., 2019; Kumar et al., 2022; Montagnac et al., 2009; Sun et al., 2017; Willett et al., 2017). Accordingly, both proteins were recently identified via unbiased proteomics as enriched in lysosomal and AV fractions from brain (Fig. 1 B) (Dumrongprechachan et al., 2022; Goldsmith et al., 2022). To assess comigration between JIP3 or JIP4 and both AVs and lysosomes in neurons, we transfected low levels of HaloTag (HT)-JIP3 or JIP4 into primary hippocampal neurons along with mScarlett (mSc)-light chain 3 (LC3)—an autophagosomal marker—and lysosome-associated membrane protein 1 (LAMP1)-mNeonGreen (mNeon). We imaged along the proximal axon, the closest 250 µm to the soma, where we previously observed the strongest impact of JIP3 siRNA on AV motility (Cason et al., 2021). In this region, the vast majority (80-100% depending on LAMP1 expression levels) of LC3 puncta colocalize with LAMP1, indicating at least one fusion event has already occurred (Cason et al., 2022; Boecker et al., 2021; Maday et al., 2012). By contrast, only about a quarter of LAMP1 puncta colocalize with LC3 (Farfel-Becker et al., 2019; Cason et al., 2022). We observed both JIP3 and JIP4 comigrating with LC3 and LAMP1 puncta (Fig. 1, C-D; S1, A-B). While high levels of JIP4 overexpression can disrupt the transport of AVs along the axon (Boecker et al., 2021), we did not observe a change in motility under the conditions tested here (Fig. S1, A-D) due to lower expression levels (see Methods for details). Likewise, overexpression of JIP3 or JIP4 did not affect LAMP1+ puncta motility, LC3 or LAMP1 density, nor colocalization between LC3 and LAMP1 in the axon (Fig. S1, A-I).

Looking specifically at the JIP3 or JIP4 puncta, we noticed that almost all of the motile puncta (moving ≥ 10µm) were directed retrograde towards the soma (Fig. 1, E-H). Because microtubules in the axon are uniformly polarized with their plus-ends out towards the axon tip and their minus ends pointing towards the soma, the minus-end-directed motor dynein is responsible for retrograde transport (Schroer et al., 1989; Schnapp and Reese, 1989; Heidemann et al., 1981). By contrast, plus-end-directed motors including kinesin-1 are responsible for anterograde transport away from the soma (Vale et al., 1985b; a). Because the vast majority of JIP3 or JIP4 puncta moved retrograde, we therefore asked whether we could observe complex formation between JIP3 or JIP4 and dynein in the axon using proximity ligation assays (PLA). PLA capitalizes on oligonucleotide complementation to identify and label proteins within 40 nm of one another in cells (Fig. 1 I) (Alam, 2018). We were indeed able to detect endogenous JIP3 or JIP4 closely apposed to endogenous dynein intermediate chain (DIC) in the axon (Fig. S1, L-M).

In live cells, HT-JIP3 or HT-JIP4 colocalized mainly with puncta positive for both LC3 and LAMP1, which can be referred to as mature AVs or autolysosomes. We therefore expressed low levels of HT-LC3 then fixed the cells and used antibodies to detect endogenous LAMP1 and assessed colocalization between LC3, LAMP1, and PLA puncta. We observed a striking colocalization between JIP3- or JIP4-DIC PLA puncta and puncta positive for both LC3 and LAMP1, with much less colocalization between PLA puncta and LC3 only or LAMP1 only puncta (Fig. 1, J-M). This finding is consistent with our previous discovery that JIP3 knockdown specifically affected the motility of mature AVs, as compared with other axonal AVs (Cason et al., 2021). Thus, we conclude that JIP3 and JIP4 complex with dynein on mature autolysosomes.

### JIP3 and JIP4 activate dynein *in vitro*

Given that JIP3 and JIP4 can each bind both dynein-dynactin and kinesin-1 (Fig. 1 A), we found it surprising that almost all of the JIP3 or JIP4 puncta in axons moved in the retrograde direction (Fig. 1 H). We therefore asked whether JIP3 or JIP4 preferentially activates dynein or kinesin motors using an *in vitro* lysate-based assay with cellular extracts prepared from COS-7 cells (Fig. 2 A). We performed all assays on dynamically growing microtubules; this allowed us to readily differentiate between the faster growing plus-end (Fig. 2 B) and the slower growing minus-end. As positive controls for dynein and kinesin activity respectively, we used BICD2^1-572^ (BICD2N), a truncated form of the known dynein activating adaptor BICD2 that lacks the autoinhibitory domain (Fig. 2, D-F); and KIF5C^1-560^ (K560), the constitutively active truncated version of KIF5C (Fig. S2, A-C). We also co-expressed HA-LIS1 to maximize the assembly of dynein complexes (Elshenawy et al., 2020; Fenton et al., 2021; Htet et al., 2020; Marzo et al., 2020). Note that we used full-length HT-JIP3 or JIP4 in our assays.

**Figure 2.**
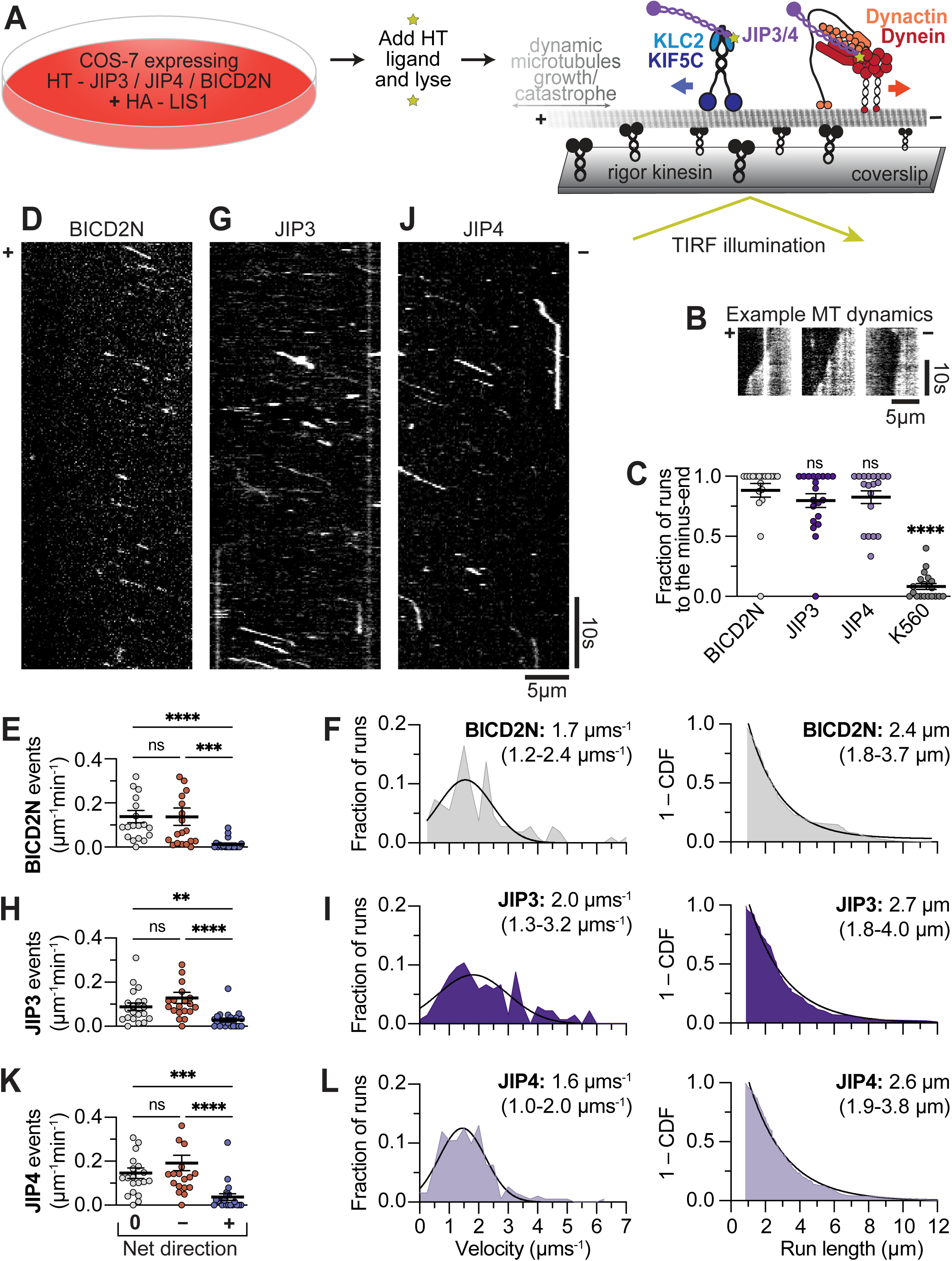
JIP3 and JIP4 induce dynein activity *in vitro*. **(A)** Schematic illustrating our single-molecule motility assay. **(B)** Example kymographs showing the growth and catastrophe dynamics used to differentiate the plus-end of the microtubule from the more stable minus-end. **(C)** Quantification of the directionality of runs on each microtubule. Runs were defined as events ≥ 0.8 µm in length towards either the minus- or plus-end of the microtubule (MT). Symbols indicate comparison to the BICD2N dynein positive control. Kruskal-Wallis test with Dunn’s multiple comparisons. n = 20 MT each. K560 v. BICD2N, P < 0.0001; K560 v. JIP3, P < 0.0001; K560 v. JIP4, P < 0.0001; BICD2N v. JIP3, P > 0.9999; BICD2N v. JIP4, P > 0.9999; JIP3 v. JIP4, P > 0.9999. **(D-F)** Example kymograph and quantification showing the activity of BICD2N-containing dynein complexes. **(G-I)** Example kymograph and quantification showing the activity of JIP3-containing dynein complexes. **(J-L)** Example kymograph and quantification showing the activity of JIP4-containing dynein complexes. All velocity histograms were fit to a Gaussian curve and all run length histograms (1– cumulative distribution frequency) were fit to a one phase decay. Listed values are median (25^th^ percentile-75^th^ percentile). n = 97-192 events. Complexes with a net direction of “0” were stationary landing events, while complexes with a net direction of “–” or “+” moved ≥ 0.8 µm towards the minus- or plus-end of the microtubule respectively. n = 20 MT each; Kruskal-Wallis test with Dunn’s multiple comparisons; JIP3 (0 v. –, P = 0.6412; 0 v. +, P = 0.0051; – v. +, P < 0.0001); JIP4 (0 v. –, P > 0.9999; 0 v. +, P = 0.0004; – v. +, P < 0.0001).

In the presence of 1mM ATP at a physiological temperature of 37°C, the majority of runs by HT-JIP3- or HT-JIP4-containing motor complexes were towards the minus-end of the microtubule (∼90%, Fig. 2 C), with velocities (∼2µm/s) and run lengths (∼3.2µm) very similar to that of BICD2N (Fig. 2, D-L). The total number of events—both runs (≥ 0.8µm net displacement) and stationary landing events (≤ 0.8 s duration with < 0.8µm net displacement)—was also similar among BICD2N (Fig. S2 B), JIP3 (Fig. 2 K), and JIP4 (Fig. 2, E, H, K). Our buffer conditions were sufficient to produce kinesin activity, as assessed using the positive control K560 (Fig. S2, A-C), yet plus-end-directed runs were rare. The few plus-end-directed events we did observe for JIP3-and JIP4-containing complexes moved slightly faster or with shorter run lengths than K560, respectively; however the low *n* for these observations prevents a direct comparison (Fig. S2, C-E).

We were surprised by the low number of plus-end-directed runs, as previous studies have reported kinesin-1 activation by JIP3 and its homolog Sunday driver (Sun et al., 2011; Watt et al., 2015). We tested whether omission of HA-LIS1 would increase the frequency of runs moving toward the microtubule plus-end, but we did not observe more kinesin runs under these conditions (Fig. S2 E). Based upon previous work, we tried combining lysate from cells expressing HT-JIP3 or JIP4 with cells expressing full-length KIF5C-HT (labelled in a different color) and GFP-KLC2. However, these experiments also failed to show substantial transport towards the plus-end of the microtubule (Fig. S2 F). One possible explanation is that previous work did not use polarity-marked microtubules (Sun et al., 2011; Watt et al., 2015); thus, it is possible that the motility they detected was in fact minus-end-directed and dynein-driven. Based upon our data, and consistent with the recent reports on the activation of dynein by JIP3 (Rao et al., 2022; Singh et al., 2022) we conclude that JIP3 and JIP4 robustly activate dynein motility, with only marginal activation of kinesin under the conditions tested. Consistent with these observations, labeled JIP3 and JIP4 both move almost exclusively retrograde in neuronal axons (Fig. 1 H).

### RAB10 overexpression differentially affects the transport of AVs and lysosomes

Many RAB GTPases have been shown to regulate motor complexes at organellar membranes (Guo et al., 2016; Amaya et al., 2016; Horgan et al., 2010; Johansson et al., 2007). In particular, RAB10 was detected in proteomics from brain-derived AVs and lysosomes and is a known JIP3/4 interactor (Waschbüsch et al., 2020; Dumrongprechachan et al., 2022; Goldsmith et al., 2022). To validate this finding, we blotted for RAB10 in fractions from total brain lysate, isolated AVs, and the AV fraction treated with Proteinase K (PK) to digest proteins specifically bound to the outer membrane as described in Goldsmith et al. (2022). RAB10 was not highly enriched in the AV fraction and, additionally, there was no significant difference between RAB10 inside (PK-protected) and outside the AV (Fig. 3 A).

**Figure 3.**
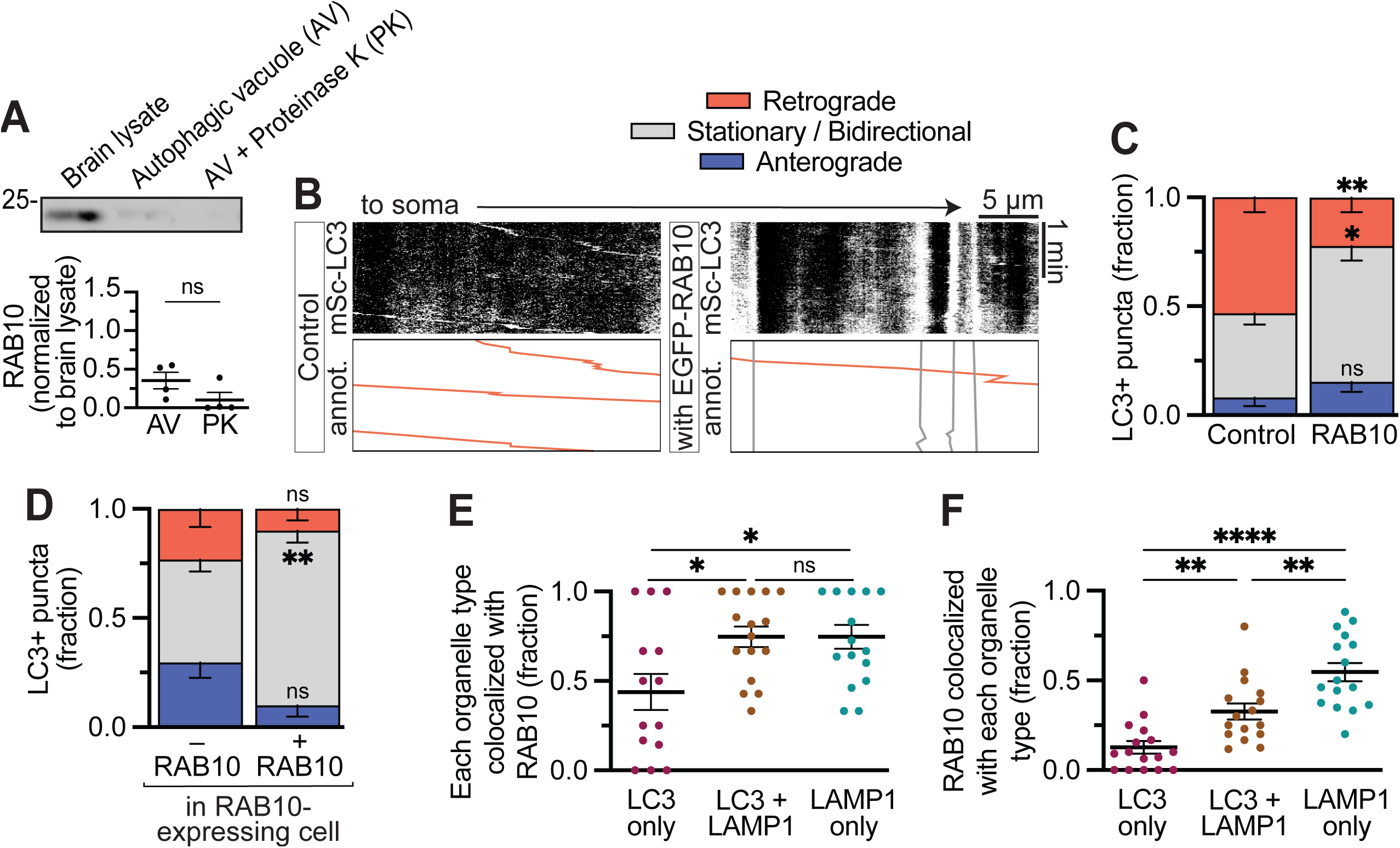
RAB10 overexpression inhibits retrograde autophagosomal transport. **(A)** Example western blot and quantification showing RAB10 in the AV fraction. n = 4 preparations; unpaired t test, P = 0.1308. **(B-C)** Example kymograph and quantification showing the fraction of LC3 moving retrograde, anterograde, or exhibiting bidirectional/stationary motion in the presence of EGFP alone (Control) or EGFP-RAB10. n = 9 neurons; two-way ANOVA with Sidak’s multiple comparisons test (anterograde, P = 0.7646; stationary/bidirectional, P = 0.0166; retrograde, P = 0.0013). **(D)** Within the RAB10-expressing cells, motility of the LC3+ puncta either colocalized with RAB10 (+ RAB10) or not (– RAB10). n = 9 neurons; two-way ANOVA with Sidak’s multiple comparisons test (anterograde, P = 0.0897; stationary/bidirectional, P = 0.0016; retrograde, P = 0.3700). **(E)** Fraction of autophagosomes (HT-LC3 only), autolysosomes (LC3+ LAMP1), or lysosomes (endogenous LAMP1 only) in fixed cells colocalized with EGFP-RAB10. n = 14-16 neurons; one-way ANOVA with Tukey’s multiple comparisons test (LC3 v. LC3 + LAMP1, P = 0.0181; LC3 v. LAMP1, P = 0.0179; LAMP1 v. LC3 + LAMP1, P > 0.9999). **(F)** Of the EGFP-RAB10 that was colocalized with LC3 and/or LAMP1, fraction colocalized with each organelle type. n = 14-16 neurons; one-way ANOVA with Tukey’s multiple comparisons test (LC3 v. LC3 + LAMP1, P = 0.0070; LC3 v. LAMP1, P < 0.0001; LAMP1 v. LC3 + LAMP1, P = 0.0030).

Surprisingly, given this lack of enrichment of RAB10 on AVs, when we expressed EGFP-RAB10 in hippocampal neurons we noted a potent dominant negative effect on LC3 motility, with EGFP-RAB10 expression leading to significantly more stationary/bidirectional AVs as compared with EGFP-tag alone (Tag; Fig. 3, B-C). In the RAB10-expressing condition, examination of RAB10 colocalization revealed that the RAB10+ AVs were almost exclusively stationary, while the few motile AVs did not colocalize with RAB10 (Fig. 3 D). RAB10 expression did not affect LC3 puncta density or colocalization with LAMP1 (Fig. S3, A-B).

Because the majority (∼85%) of the AVs in the proximal axon have fused with a lysosome and are LAMP1+ (Fig. S3 B), we assessed colocalization between EGFP-RAB10, HT-LC3, and endogenous LAMP1 in fixed neurons. We found that RAB10 predominantly colocalized with LAMP1+ organelles, including both autolysosomes and lysosomes (Fig. 3, E-F). Hence, we asked whether LAMP1+ puncta motility is also affected by RAB10 expression. There was no gross effect on LAMP1 puncta motility (Fig. 4, A-B). However, when we specifically quantified the motile puncta (moving ≤ 10µm in either direction during a 2 min video), we found a significant shift from a mild retrograde bias (∼60%) in the GFP expressing cells to a mild anterograde bias (∼57%) in the RAB10 expressing cells (Fig. 4 C). Oddly, however, when we examined the LAMP1 puncta colocalized with RAB10, we found the majority to be stationary/bidirectional (Fig. 4 D), like for LC3 (Fig. 3 D). RAB10 expression had no effect on LAMP1 colocalization with LC3, but was associated with a mild decrease in LAMP1 puncta in the axon (Fig. S3, C-D).

**Figure 4.**
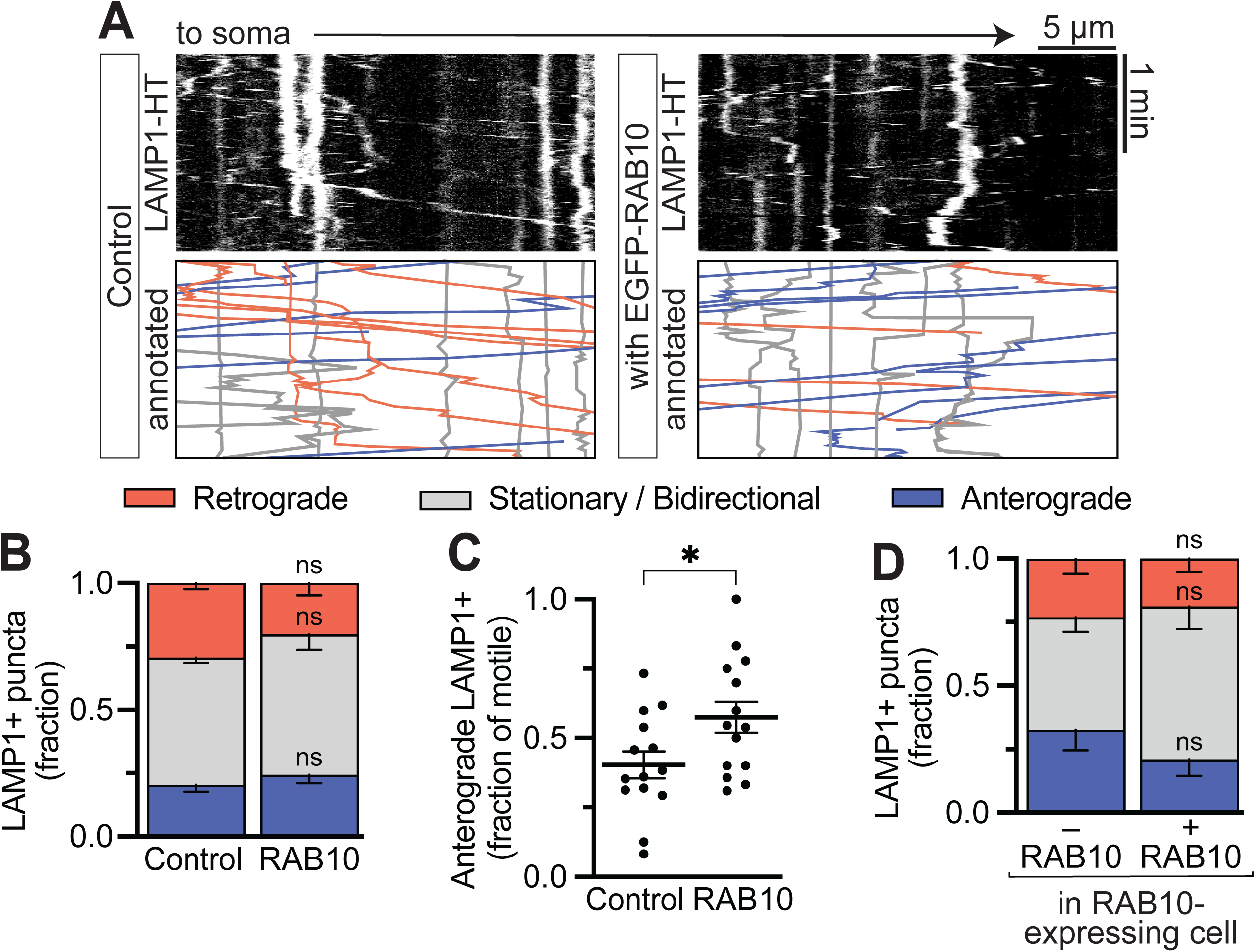
RAB10 overexpression shifts the lysosomal population towards anterograde motility. **(A-B)** Quantification and example kymographs showing the fraction of LC3 moving retrograde, anterograde, or exhibiting bidirectional/stationary motion in the presence of EGFP alone (Control) or EGFP-RAB10. n = 14 neurons; two-way ANOVA with Sidak’s multiple comparisons test (anterograde, P = 0.8445; stationary/bidirectional, P = 0.7328; retrograde, P = 0.2696). **(C)** Fraction of the motile LAMP1 puncta (moving ≤ 10µm in either direction during a 2 min video) moving anterograde. n = 14 neurons; unpaired t test (P = 0.0297). **(D)** Within the RAB10-expressing cells, motility of the LAMP1+ puncta either colocalized with RAB10 (+ RAB10) or not (– RAB10). n = 9 neurons; two-way ANOVA with Sidak’s multiple comparisons test (anterograde, P = 0.5595; stationary/bidirectional, P = 0.3011; retrograde, P = 0.9651).

We used PLA (Fig. 1 I) to determine whether the changes in motility were due to a disruption in the formation of JIP3/4-dynein complexes. We found that the number of JIP3/4-DIC PLA puncta did not change upon RAB10 expression (Fig. S3, E-F), nor did the colocalization between JIP3/4-DIC PLA puncta and autophagosomes, autolysosomes, or lysosomes (Fig. S3, G-H). Hence, the decrease in AV motility and the increase in LAMP1 anterograde motility is not due to loss of JIP3/4-dynein complexes. Together, these results suggest that RAB10 can modulate the motility of AVs and lysosomes, potentially by affecting kinesin recruitment and/or activation, rather than by decreasing dynein recruitment or activation.

### ARF6 GDP-locked mutant disrupts autophagosomal transport

To better understand the regulation of JIP3/4-dependent AV motility, we probed the proteomic data for other possible small GTPases that might affect motor activity or motor coordination. We identified the candidate ARF6, which is present in both the AV and lysosomal proteomic datasets (Dumrongprechachan et al., 2022; Goldsmith et al., 2022) and whose enrichment on the outside of the AV we could validate via immunoblot (Fig. 5 A). ARF6 is a known JIP3/4 interactor and has been previously shown to modulate JIP3/4 motor binding based upon its GTP-binding state: ARF6-GTP increases the binding between JIP3/4 and the dynactin subunit p150^Glued^ (p150) while ARF6-GDP increases binding between JIP3/4 and kinesin light chain (KLC; Fig. 5 B) (Montagnac et al., 2009). Hence, we transfected CFP-ARF6 GTP-locked (Q67L) and GDP-locked (T27N) mutants into our primary hippocampal neurons and assessed the motility of LC3 puncta. While expression of wildtype (ARF6^WT^) and ARF6^QL^ did not have an obvious effect on LC3 motility, ARF6^TN^ expression induced a robust loss of AV motility and a significant increase in the pausing of LC3 puncta (Fig. 5, C-E). However, there was no effect on either LC3 puncta density or AV maturation, measured either by LAMP1 colocalization (Fig. S4, A-B).

**Figure 5.**
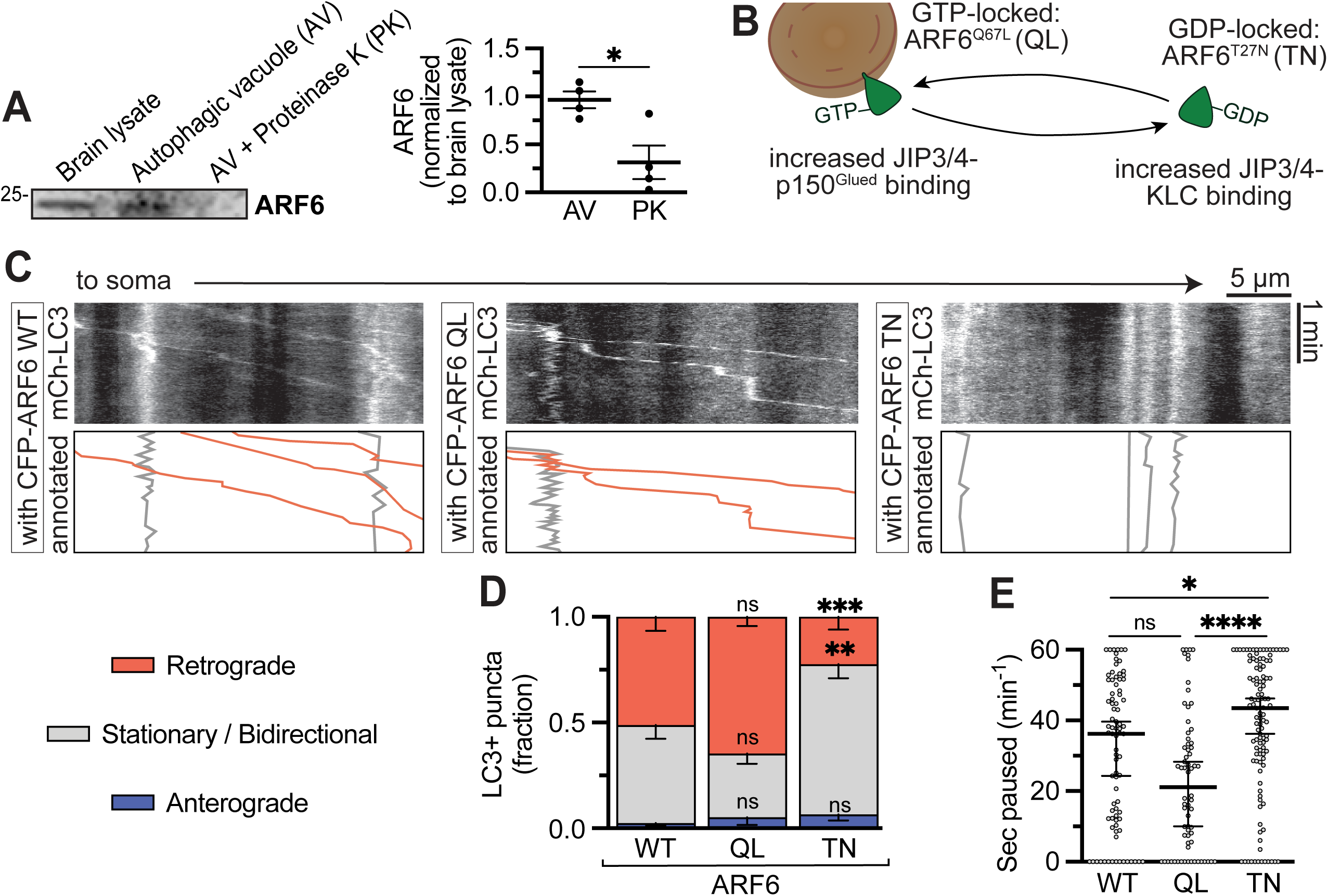
ARF6 regulates the motility of autophagosomes in the axon in a GTP-dependent fashion. **(A)** Example western blot and quantification showing ARF6 in the AV fraction. n = 4 preparations; unpaired t test, P = 0.0159. **(B)** Schematic illustrating the general characteristics of GTP- or GDP-ARF6 and the locked point mutants. **(C-D)** Example kymographs and quantification of mCherry (mCh)-LC3 motile fractions under the expression of CFP-ARF6^WT^, ARF6^Q67L^, or ARF6^T27N^. n = 15-18 neurons; two-way ANOVA with Tukey’s multiple comparisons; anterograde (WT v. QL, P = 0.9333; WT v. TN, P = 0.8377; QL v. TN, P = 0.9795); stationary/bidirectional (WT v. QL, P = 0.0824; WT v. TN, P = 0.0021; QL v. TN, P < 0.0001); retrograde (WT v. QL, P = 0.1723; WT v. TN, P = 0.0003; QL v. TN, P < 0.0001). Symbols indicate comparison to ARF6^WT^. **(E)** Number of seconds paused per min in each of the three conditions (for all AVs). n = 83-111 puncta; Kruskal-Wallis test with Dunn’s multiple comparisons; WT v. QL, P = 0.1526; WT v. TN, P = 0.0327; QL v. TN, P < 0.0001.

### ARF6 GTP-locked mutant decreases retrograde lysosome pausing

We next assessed motility of LAMP1 puncta in the axon upon ARF6 expression. Here, we observed a very different effect. While gross LAMP1 motility was not significantly different in neurons expressing ARF6^WT^ or ARF6^TN^ expression, expression of the GTP-locked ARF6^QL^ mutant led to a decrease in the stationary/bidirectional fraction of LAMP1 puncta (Fig. 6, A-B). Expression of either locked mutant led to decreased pausing time, although the effect was much bigger in ARF6^QL^-expressing cells (Fig. 6 C). We evaluated pausing time for LAMP1 puncta colocalized with LC3 (autolysosomes) and those not colocalized (lysosomes) and saw that both organelle subgroups were affected by ARF6^QL^ expression (Fig. 6 D). Neither the LAMP1 puncta colocalized with LC3 nor the LAMP1 density was affected (Fig. S4 D-E). However, when we split the LAMP1 puncta into motile fractions, we saw that the pausing effect was limited to the retrograde LAMP1 fraction, with no significant effect on anterograde-moving LAMP1 puncta (Fig. 6, E-F). Thus, GTP-bound but not GDP-bound ARF6 increases the efficiency of the dynein complex on LAMP1-positive organelles.

**Figure 6.**
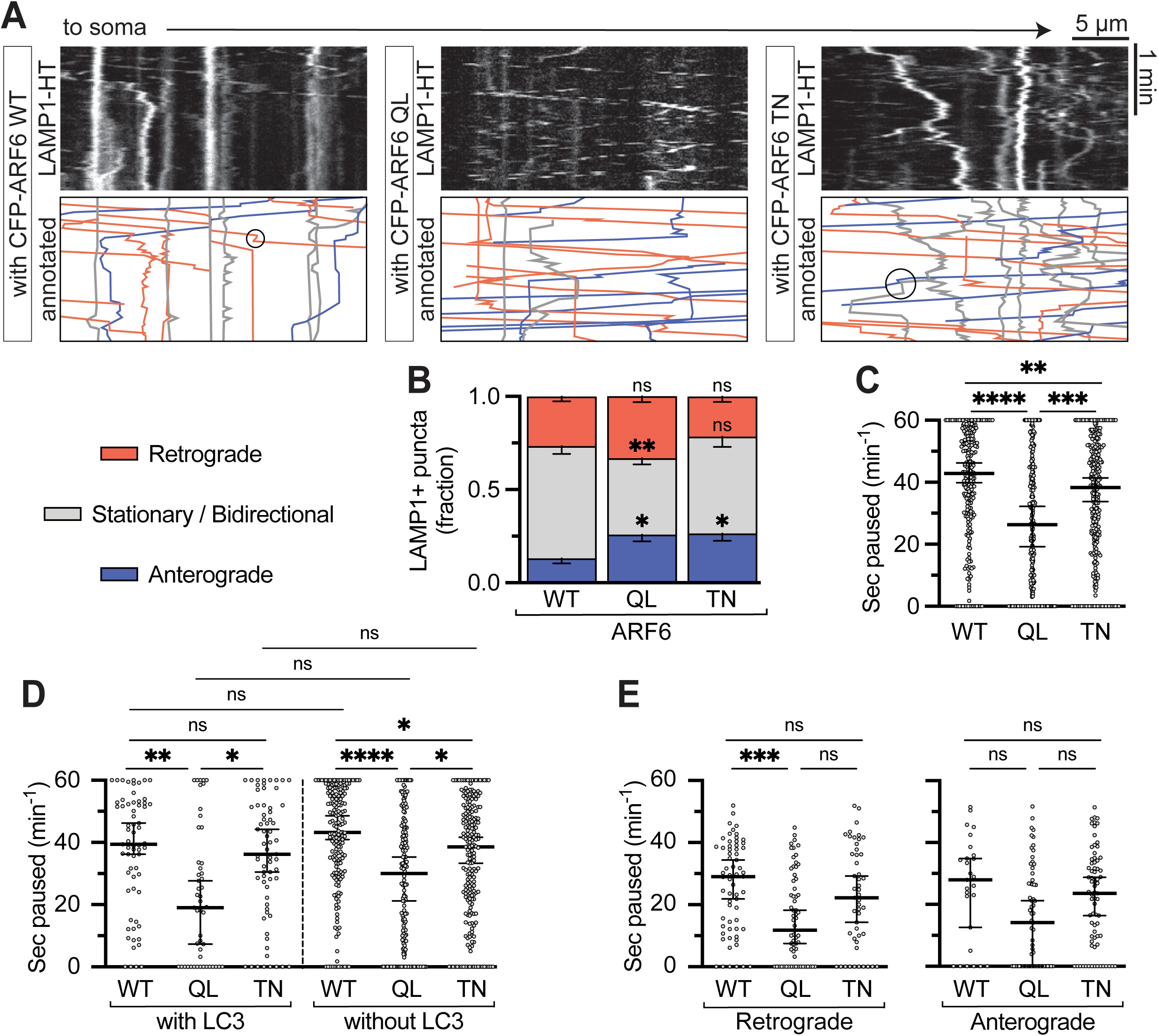
ARF6 GTP-locked mutant decreases retrograde lysosome pausing. **(A-B)** Example kymographs and quantification of LAMP1-HT motile fractions under the expression of CFP-ARF6^WT^, ARF6^Q67L^, or ARF6^T27N^. n = 10 neurons; two-way ANOVA with Tukey’s multiple comparisons; anterograde (WT v. QL, P = 0.0428; WT v. TN, P = 0.0314; QL v. TN, P = 0.9915); stationary/bidirectional (WT v. QL, P = 0.0011; WT v. TN, P = 0.2466; QL v. TN, P = 0.0960); retrograde (WT v. QL, P = 0.4213; WT v. TN, P = 0.6013; QL v. TN, P = 0.0729). Symbols indicate comparison to ARF6^WT^. **(C-F)** Number of seconds paused per min in each of the 3 conditions for all LAMP1 puncta (C; n = 223-274 puncta; WT v. QL, P < 0.0001; WT v. TN, P = 0.0059; QL v. TN, P = 0.0001), LAMP1 with (D; n = 49-67 puncta; WT v. QL, P = 0.0029; WT v. TN, P > 0.9999; QL v. TN, P = 0.0265) and without LC3 (D; n = 174-206 puncta; WT v. QL, P < 0.0001; WT v. TN, P = 0.0198; QL v. TN, P = 0.0431), and all the retrograde-(E; n = 49-65 puncta; WT v. QL, P = 0.0004; WT v. TN, P = 0.6390; QL v. TN, P = 0.0574) or anterograde-(E; n = 27-73 puncta; WT v. QL, P = 0.0507; WT v. TN, P > 0.9999; QL v. TN, P = 0.0846) moving LAMP1 puncta. Kruskal-Wallis test with Dunn’s multiple comparisons.

### ARF6 GTPase status is locally regulated by GAPs and GEFs on the membrane

We next asked whether wildtype ARF6 may be converted locally at AVs and lysosomes into a GTP-bound and GDP-bound state respectively. The nucleotide state of small GTPases is regulated by GTPase activating proteins (GAPs), which induce GTP-to-GDP hydrolysis, and guanine exchange factors (GEFs), which induce release of GDP and binding of a new GTP molecule (Fig. 7 A). There are 10 known ARF6 GEFs and 20 GAPs. 60% of ARF6 GEFs and 60% of ARF6 GAPs were detected in lysosomal proteomics (Fig. 7 B) (Dumrongprechachan et al., 2022). Because LC3+ puncta in cells expressing ARF6^QL^ behaved similarly to wildtype-expressing cells, we hypothesized that AVs would be enriched for ARF6 GEFs. Indeed, 50% of ARF6 GEFs and only 20% of ARF6 GAPs were detected in AV proteomics (Fig. 7 B) (Goldsmith et al., 2022). We used immunoblotting to validate the GAPs and GEFs detected in an AV-enriched fraction from brain, and found that the GEFs were highly enriched in the AV fraction and significantly localized to the outer membrane of the AV as judged by protease sensitivity (Fig. 7, C-G). By comparison, the ARF6 GAPs were less enriched and not significantly localized to the outer membrane (Fig. 7, H-J). These observations support a model in which ARF6 GEFs are localize to the AV membrane to locally enrich for ARF6-GTP, which can in turn can recruit and/or activate JIP3/4-containing dynein complexes.

**Figure 7.**
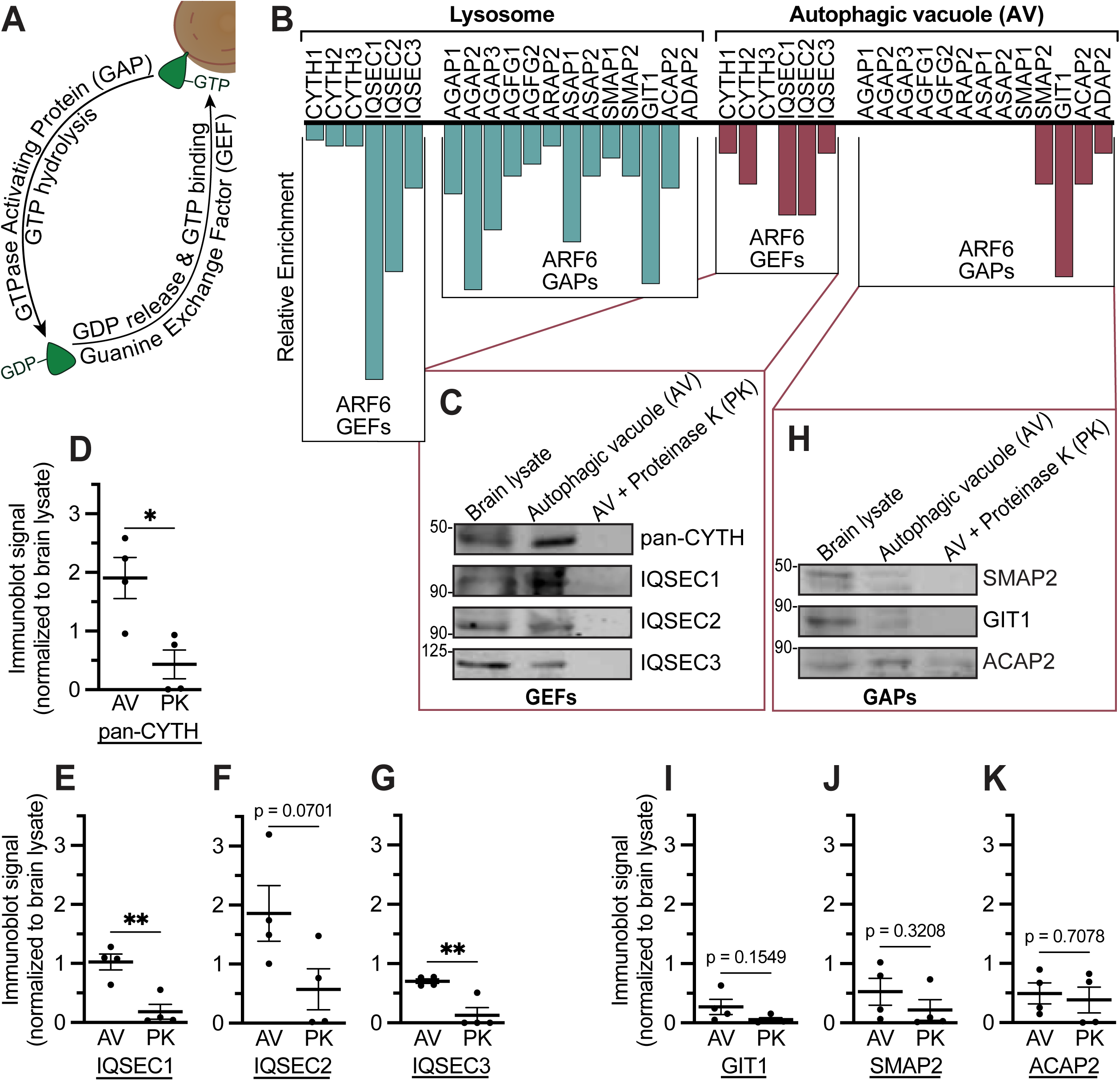
ARF6 GEFs are enriched on the outer AV membrane and may act to locally enrich ARF6-GTP. **(A)** Schematic illustrating the GAP-GEF cycle for small GTPases. **(B)** Relative enrichment (normalized number of peptides, see Methods for details) for ARF6 GEFs and GAPs in the proteomics performed by Goldsmith et al., (2022) and Dumrongprechachan et al., (2022). All of the known ARF6 GEFs and GAPs that were detected in either organelle enrichment are listed in the figure. Note that some ARF6 GAPs/GEFs were not found in either enrichment [EFA6A-D (GEFs), GIT2 (GAP), ADAP1 (GAP), ACAP1,3 (GAP), ASAP3 (GAP), ARAP1,3 (GAP)]. **(C)** Example western blot and **(D-G)** quantification showing GEF enrichment in the AV fraction. **(H)** Example western blot and **(I-K)** quantification showing GAPs in the AV fraction. n = 4 preparations; unpaired t test; (D) P = 0.0139; (E) P = 0.0038; (F) P = 0.0701; (G) P = 0.0050; (I) P = 0.1549; (J) P = 0.3208; (K) P = 0.7078.

### ARF6 increases the interaction of the JIP3/4-dynein complex with microtubules

Finally, we asked how ARF6 might affect the behavior of JIP3/4-containing motor complexes using *in vitro* motility assays. We found that overexpression of GTP-locked ARF6 significantly increased the microtubule landing events of JIP3- or JIP4-containing motor complexes (Fig. 8, A-C), although the frequency of plus- and minus-end-directed events did not change (Fig. 8, D-E). Interestingly, ARF6^TN^ induced the same effect, a significant increase in microtubule-binding events (Fig. S5, A-E). There was no effect on the velocity of motile complexes in either ARF6^QL^ or ARF6^TN^ conditions, but there was a mild reduction in run length upon ARF6 inclusion (∼28%; Fig. S5, F-G).

**Figure 8.**
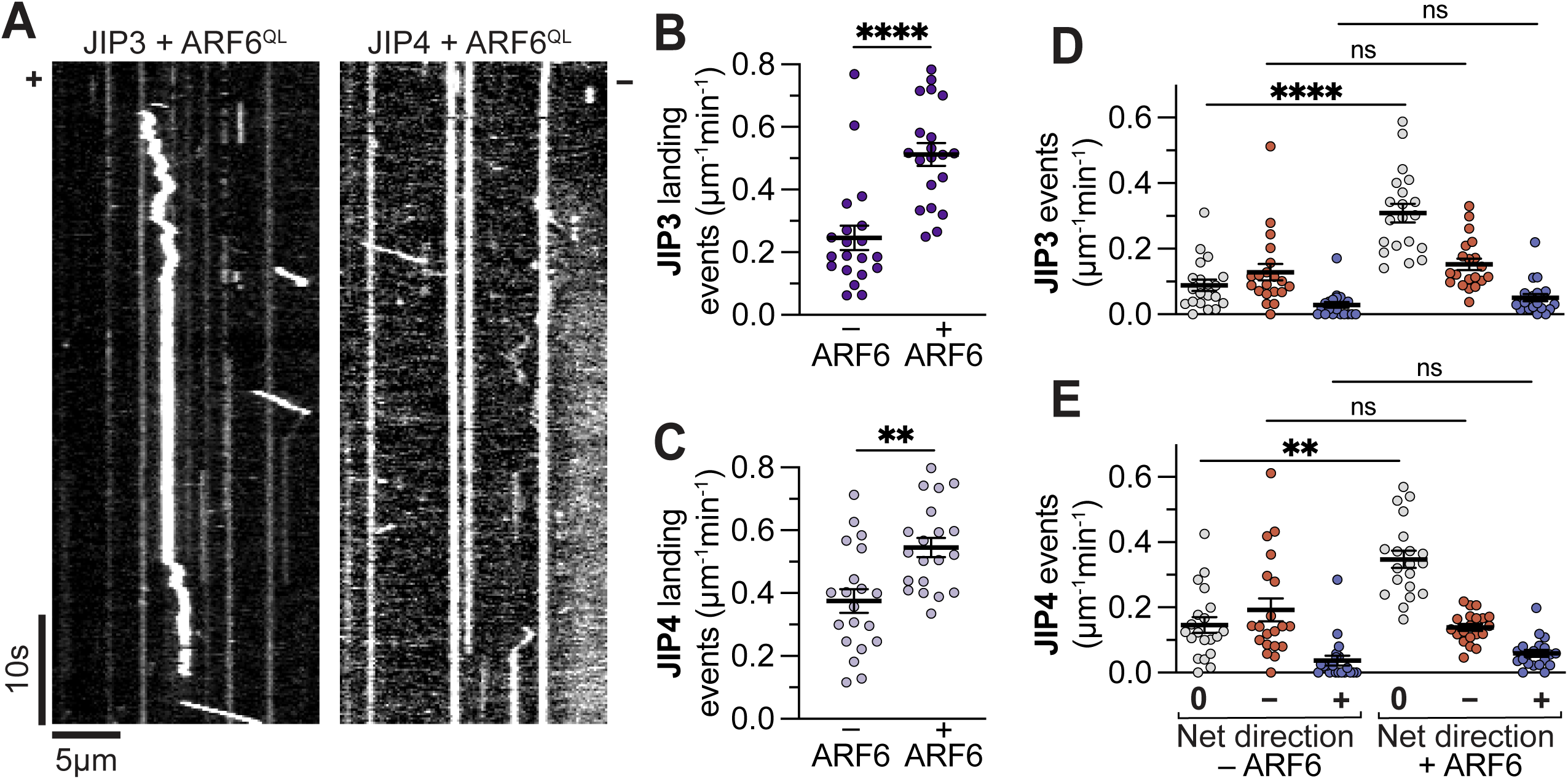
ARF6 induces the recruitment of JIP3/4 to microtubules. **(A)** Example kymographs showing the activity of JIP3- and JIP4-containing complexes in the presence of CFP-ARF6^Q67L^. **(B-C)** Quantification of the number of total landing events for JIP3- and JIP4-containing complexes in the presence or absence of CFP-Arf6^Q67L^. n = 20 MT each; unpaired t test; JIP3, P < 0.0001; JIP4, P = 0.0011. **(D-E)** Number of events (per µm microtubule per min) observed for JIP3-containing and JIP4-containing complexes, either in the presence or absence of ARF6^Q67L^. Complexes with a net direction of “0” were stationary landing events, while complexes with a net direction of “–” or “+” moved ≥ 0.8 µm towards the minus- or plus-end of the microtubule respectively. Note that the –ARF6 data from is repeated from Figure 2. Kruskal-Wallis test with Dunn’s multiple comparisons. n = 20 MT each. JIP3 –ARF6 v. + ARF6: 0, P < 0.0001; –, P > 0.9999; +, P > 0.9999. JIP4 –ARF6 v. + ARF6: 0, P = 0.0033; –, P > 0.9999; +, P > 0.9999.

Because JIP3 and JIP4 do not directly bind microtubules, the increased landing events must be due to increased interaction with a microtubule-binding protein. The dynactin subunit p150 interacts directly with JIP3/4 and with microtubules via its CAP-Gly domain; additionally, ARF6 binding modulates the interaction between p150 and JIP3/4 (Peris et al., 2006; Ayloo et al., 2014; Moughamian and Holzbaur, 2012; Montagnac et al., 2009). Finally, p150 binding to the microtubule in the absence of dynein induces statically bound and/or diffusive behaviors (Ayloo et al., 2014; Feng et al., 2020), similar to what we observe. Therefore, we propose that ARF6 increases the efficiency of JIP3/4-containing motor complexes in cells by increasing the microtubule association of JIP3/4 through p150.

## Discussion

Here, we demonstrate that two related scaffolding proteins, JIP3 and JIP4, both activate dynein in vitro, and form a complex with dynein on mature AVs (autolysosomes) in neuronal axons (Fig. 1, 2). We identify two small GTPases that interact with JIP3/4 and affect the axonal transport of AVs and other LAMP1+ organelles. RAB10 overexpression halts the retrograde transit of AVs and increases the anterograde bias observed for LAMP1-positive puncta in the axon (Fig. 3, 4). ARF6 also regulates AV motility, in a GTP-binding-dependent fashion: GTP-locked ARF6 decreases the pausing of retrograde-moving LAMP1 puncta, while GDP-locked ARF6 increases the fraction of stationary AVs (Fig. 5, 6). Further, ARF6 GEFs are enriched on the outer AV membrane (Fig. 7), meaning ARF6-GTP can be locally upregulated. We propose that locally generated ARF6-GTP recruits the JIP3/4-dynein-dynactin complex and also enhances the association of the complex with microtubules (Fig. 8), leading to more efficient transport of AVs toward the soma.

Concurrent with our study, two other groups have shown that purified recombinant truncated JIP3 is sufficient to activate dynein-mediated motility *in vitro* (Rao et al., 2022; Singh et al., 2022). Our study adds to this growing body of work, as the approaches used here (1) include endogenous binding partners, which negates the need to use truncated constructs; (2) is performed at physiological temperature on dynamically growing microtubules; and (3) includes competing kinesin complexes. Our average JIP3 velocity (∼1.8 µms^-1^) is higher than that observed by the other groups [0.7 µms^-1^ (Rao et al., 2022); 1 µms^-1^ (Singh et al., 2022)], most likely due to the more physiological assay temperature (37°C vs. room temperature). It is, however, consistent with previous observations of dynein activation using lysate assays performed at 37° (Fenton et al., 2021).

Previous work has reported that the Drosophila JIP3 ortholog Sunday driver (syd) activates kinesin-1 (Sun et al., 2011). However, these assays were performed using mammalian cell lysate at room temperature on stabilized microtubules without polarity labelling, and the average velocities (0.6-1.0 µms^-1^) and run lengths (3-5.5 µm) observed suggest that minus end-directed motility may have dominated in their assays, as these values are more consistent with dynein-mediated transport (Olenick et al., 2016; Urnavicius et al., 2018; Fenton et al., 2021; Canty et al., 2021; Fu et al., 2014; Fu and Holzbaur, 2013; Rao et al., 2022; Singh et al., 2022; Sun et al., 2011). Similar assays performed with mammalian JIP3 (Watt et al., 2015) resulted in velocities (∼0.25 µms^-1^) more consistent with kinesin-1 activation; however, the relatively short run lengths (∼0.75 µm) suggest that JIP3 may require additional effectors to fully activate kinesin-1 motility. Consistent with this conclusion, binding assays suggest that JIP3 and the unrelated motor effector protein JNK-interacting protein 1 (JIP1) cooperatively activate kinesin-1 (Sun et al., 2017). JIP1 alone is unlikely to be sufficient for kinesin-1 activation (Blasius et al., 2007; Sun et al., 2017), but in single molecule assays using cell lysates, JIP1 overexpression increases the number of kinesin-1-driven motility events (Fu and Holzbaur, 2013; Fu et al., 2014). Binding assays suggest that the binding of JIP1 to KHC and KLC, concurrent with the binding of JIP3 to KLC, is necessary to fully relieve kinesin-1 autoinhibition (Sun et al., 2017). Further, overexpression of either JIP1 or JIP3 leads to the accumulation of the other adaptor at microtubule plus-ends in cells, suggesting cotransport with kinesin (Hammond et al., 2008). While JIP3 does not oligomerize with JIP4, JIP1 and JIP3 interact both directly and indirectly through KLC, where they bind distinct residues in the tetratricopeptide repeat (TPR) domain (Hammond et al., 2008; Kelkar et al., 2005). Of note, JIP1 and JIP3 have been implicated in the anterograde transport of many of the same organelles, including synaptic vesicle proteins, TrkB receptor, amyloid precursor protein (APP), mitochondria, and signaling proteins such as JNK (Choudhary et al., 2017; Horiuchi et al., 2005; Sun et al., 2017; Fu and Holzbaur, 2013; Drerup and Nechiporuk, 2013; Sato et al., 2015), further supporting a cooperative interaction between these motor activators.

Notably, both JIP1 and JIP3/4 have been previously implicated in the transport of RAB10+ vesicles (Kluss et al., 2022; Bonet-Ponce et al., 2020; Deng et al., 2014). In our study, RAB10 appears to increase the recruitment and/or activation of kinesin on LC3+ and LAMP1+ puncta. RAB10, like many other RABs, is regulated by phosphorylation with phospho-RAB10 (especially T73) being generally more active and membrane-associated (Yan et al., 2018; Lara Ordóñez et al., 2022; Kluss et al., 2022; Wauters et al., 2020; Waschbüsch et al., 2020; Homma et al., 2021). RAB10, especially phospho-RAB10, regulates the motility of multiple kinesin-1 and kinesin-3 cargoes within cells (Etoh and Fukuda, 2019; Deng et al., 2014; Taylor et al., 2015; Zajac and Horne-Badovinac, 2022). RAB10 can directly complex with kinesin-3 (KIF13) and regulate its activity (Etoh and Fukuda, 2019; Zajac and Horne-Badovinac, 2022). However, its interaction with kinesin-1 (KIF5) must be mediated by adaptor proteins. We therefore propose an integrated model (Fig. 9 A) in which JIP3 or JIP4, together with JIP1, mediate the anterograde transport of RAB10+ cargo, including some populations of lysosomes. Phospho-RAB10 recruits JIP3/4 and JIP1 to LAMP1+ organelles; upon RAB10 binding, JIP3/4 and JIP1 bind to kinesin-1 to induce the anterograde transit of the organelle (Fig. 3, 4). RAB10 can also directly bind kinesin-3 to induce anterograde transport, circumventing the JIP3/4-JIP1 complex.

**Figure 9.**
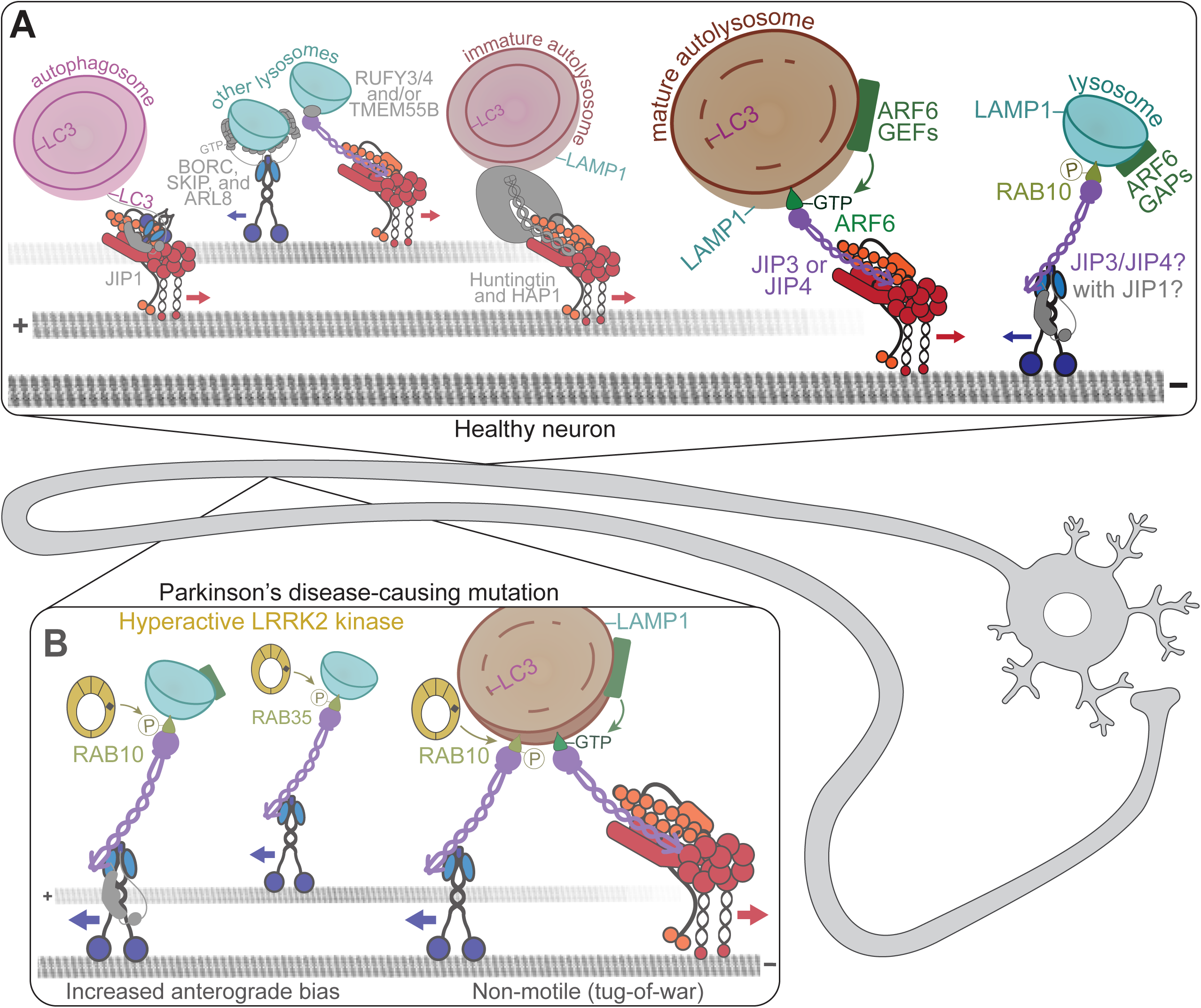
Integrated model of autophagosome, autolysosome, and lysosome transport along axons. **(A)** We demonstrate that the GTP-bound ARF6 is enriched on AV membranes, where it can recruit its interacting partners JIP3 or JIP4. JIP3/4 can then recruit dynactin and dynein and activate minus-end-directed retrograde motility especially of autolysosomes. By contrast, we propose that ARF6 is removed from the lysosomal membrane by local ARF6 GAP activity, which may be promoted by the presence of phosphorylated RABs, including RAB10. RAB10 plays an unknown role in transport, but seems to induce anterograde transit, possibly through a JIP3- or JIP4-JIP1-kinesin-1 complex. These motor complexes are not the only ones involved in AV or lysosome transport; we highlight a few complementary complexes on the left. **(B)** One mechanism by which this pathway may be disrupted in neurodegeneration is via hyperphosphorylation of RABs. The disease-causing mutations in LRRK2 kinase result in increased phospho-RABs and also increased recruitment of kinesin-1 to the AV membrane. However, the resulting loss of AV motility can be rescued by expressing GTP-locked ARF6; thus these motor-regulatory mechanisms are interconnected and possibly competitive.

RAB10 is a known target of the kinase LRRK2, which is hyperactive in some genetic forms of Parkinson’s disease (Yan et al., 2018; Wauters et al., 2020; Bonet-Ponce et al., 2020). Our group previously showed that hyperactive, disease-associated LRRK2 increased the level of phospho-RABs and kinesin present on the outer membrane of AVs (Boecker et al., 2021). Additionally, hyperactive LRRK2 increased the recruitment of JIP4 and, to a lesser degree, JIP3 to the AV membrane (Boecker et al., 2021). We therefore hypothesize that phospho-RAB10 forms a complex with JIP4 to recruit and activate kinesin (Fig. 9 B). If exogenously expressed at high levels, JIP4 can block AV motility (Boecker et al., 2021), but when expressed at more modest levels, JIP4 comigrates with autolysosomes (Fig. S2), like JIP3. It is unknown whether JIP4 can also interact with JIP1 to activate kinesin-1. However, based upon our work and others, we suggest that JIP3 and JIP4 are functionally redundant with their primary difference being expression in different tissues (Gowrishankar et al., 2021; Tuvshintugs et al., 2014; Sato et al., 2015). Thus, in non-neuronal cells where JIP3 is not expressed, JIP4 may replace JIP3 in the kinesin activation complex.

JIP3 and JIP4 can also induce retrograde transport of LAMP1+ organelles, including autolysosomes. ARF6 was previously shown to block JIP3 or JIP4 binding to kinesin, possibly through steric hindrance: the KLC and ARF6 binding sites are highly overlapping (Vilela et al., 2019; Cockburn et al., 2018; Hammond et al., 2008; Llinas et al., 2016; Isabet et al., 2009; Montagnac et al., 2009) (Fig. 1 B). Instead, ARF6 enhances the interaction between JIP3/4 and the dynactin subunit p150^Glued^ (Montagnac et al., 2009). The increased frequency of microtubule landing events for JIP3/4—in the absence of changes to other motility parameters—induced by addition of ARF6 to our *in vitro* assays (Fig. 8) is consistent with an increased interaction with p150^Glued^, which contains a microtubule-binding domain but no motor activity (Ayloo et al., 2014; Feng et al., 2020). Microtubule binding, especially through dynactin and general dynein effectors like CLIP-170, has previously been shown to be important for dynein recruitment and initiation of motility (Moughamian and Holzbaur, 2012; Moughamian et al., 2013; Nirschl et al., 2016; McKenney et al., 2016). Stepwise recruitment of dynactin and then dynein to the microtubule has not previously been shown for cargo-specific activating activators for dynein. However, the JIP3/4-related dynein effector Rab7-interacting protein (RILP) was previously shown to bind to dynactin prior to the initiation of dynein activity, suggesting a similar mechanism (Johansson et al., 2007). While RILP has not yet been shown to activate motor activity *in vitro*, its N-terminus—including the motor binding domains—is very similar to that of JIP3 and JIP4 (Celestino et al., 2022; Vilela et al., 2019; Matsui et al., 2012). We therefore hypothesize that the formation of an initial microtubule-bound cargo-binding protein-dynein activator-dynactin complex, such as the ARF6-JIP3/4-dynactin complex, is a common mechanism in the initiation of dynein-mediated transport of diverse cargos.

In single molecule motility assays, we observed no difference between GTP- and GDP-locked ARF6 (Fig. 8, S9). However, in neurons, expression of GTP- and GDP-locked ARF6 induced significant changes in organelle transport (Fig. 5, 6). We hypothesize that this difference may be due to a key role for bound nucleotide in regulating the membrane association of ARF6. ARF6 contains a myristoyl anchor and binds more tightly to membranes in its GTP-bound state than in its GDP-bound state (Ménétrey et al., 2000; Duellberg et al., 2021). In our *in vitro* assays, membranes are first removed via centrifugation of cell lysates, making this assay less sensitive to effects of the nucleotide state of ARF6. Therefore, we propose that the differential effects observed in cells and in vitro when comparing GTP- and GDP-locked ARF6 are likely due to increased membrane interaction and the subsequent stepwise recruitment of JIP3 or JIP4, dynactin, and dynein (Fig. 9 A).

Interestingly, overexpression of GTP-locked ARF6 is sufficient to ameliorate the AV motility phenotype observed in hyperactive LRRK2 mutant conditions (Dou et al., 2022). Under increased LRRK2 activity, JIP3/4 seem to exhibit enhanced interaction with kinesin and phosphorylated RABs (Dou et al., 2022; Boecker et al., 2021; Kluss et al., 2022; Bonet-Ponce et al., 2020). In these conditions, GTP-locked ARF6 expression presumably scaffolds the formation of more JIP3/4-dynein-dynactin complexes to compete with the RAB-JIP3/4-kinesin-1 complexes (Dou et al., 2022). ARF6 and phospho-RAB10 (or RAB35) may compete to bind the same limited pool of JIP3/4, or excess JIP3/4 may be available for recruitment into multiple motor complexes concurrently (Bonet-Ponce et al., 2020; Miyamoto et al., 2014; Kobayashi and Fukuda, 2012). While ARF6 and RAB10 bind different regions of JIP3/4, the ARF6 binding site and the KLC binding site are mutually exclusive (Isabet et al., 2009); thus it is likely that these are two completely discrete complexes.

It is possible that crosstalk occurs locally between ARF6 GAPs and GEFs and RAB kinases and phosphatases to prevent local activation of both dynein and kinesin, which would result in non-processive tug-of-war. For example, the RAB10/RAB35 effector ACAP2 is known to also function as a GAP for ARF6 (Shi and Grant, 2013; Shi et al., 2012; Kobayashi and Fukuda, 2012; Miyamoto et al., 2014). ACAP2 is present in both the AV and lysosome proteomics datasets from brain (Goldsmith et al., 2022; Dumrongprechachan et al., 2022); however, we found that ACAP2 was primarily an AV cargo, not on the outer membrane, consistent with local enrichment for ARF6-GTP on the AV membrane. However, ACAP2 enrichment on lysosomes may locally promote GTP hydrolysis by ARF6, leading to dissociation of the GTPase from the membrane. Interestingly, analysis of the RAB35-ACAP2 structure indicates that ACAP2 specifically binds to the LRRK2 phosphorylation site within RAB35 (Lin et al., 2019). Thus, LRRK2 kinase activity may locally enhance the formation of RAB-JIP3/4-kinesin-1 complexes, and also prevent local accumulation of ARF6-GTP and the resultant activation of dynein-dynactin activity on lysosomes.

Hyperphosphorylated RAB10 is induced by Parkinson’s disease-causing mutations in LRRK2, and is also a hallmark pathological feature of Alzheimer’s disease (Yan et al., 2018). Mutations in the JIP1-JIP3 cargo APP cause familial Alzheimer’s disease; further, both axonal transport and autophagy are disrupted in a multitude of neurodegenerative diseases (Wong and Holzbaur, 2015; Kins et al., 2006; Guillaud et al., 2020; Goldstein, 2012). Mutations in JIP3 cause a rare neurodevelopmental disorder (Platzer et al., 2019) and double-knockout of JIP3 and JIP4 leads to robust neurodegeneration (Sato et al., 2015; Gowrishankar et al., 2021). Additionally, ARF6 knockout in neurons leads to defects in axonal development (Akiyama et al., 2014). Because these proteins and the processes they regulate are all dysfunctional in neurodevelopmental and/or neurodegenerative disease, it is essential that we continue to tease apart the detailed mechanisms involved in order to better inform therapy development.

## Acknowledgements

This research was supported by NIH grant R35 GM126950 to E.L.F.H. The authors declare no competing financial interests. We gratefully acknowledge Mariko Tokito for her tireless cloning, Dan Dou and Alex Boecker for discussions, and Juliet Goldsmith for her generosity with autophagosome fractionation samples.

## Materials and methods

### Plasmids and reagents

Constructs, all of which were verified by DNA sequencing, include the following:

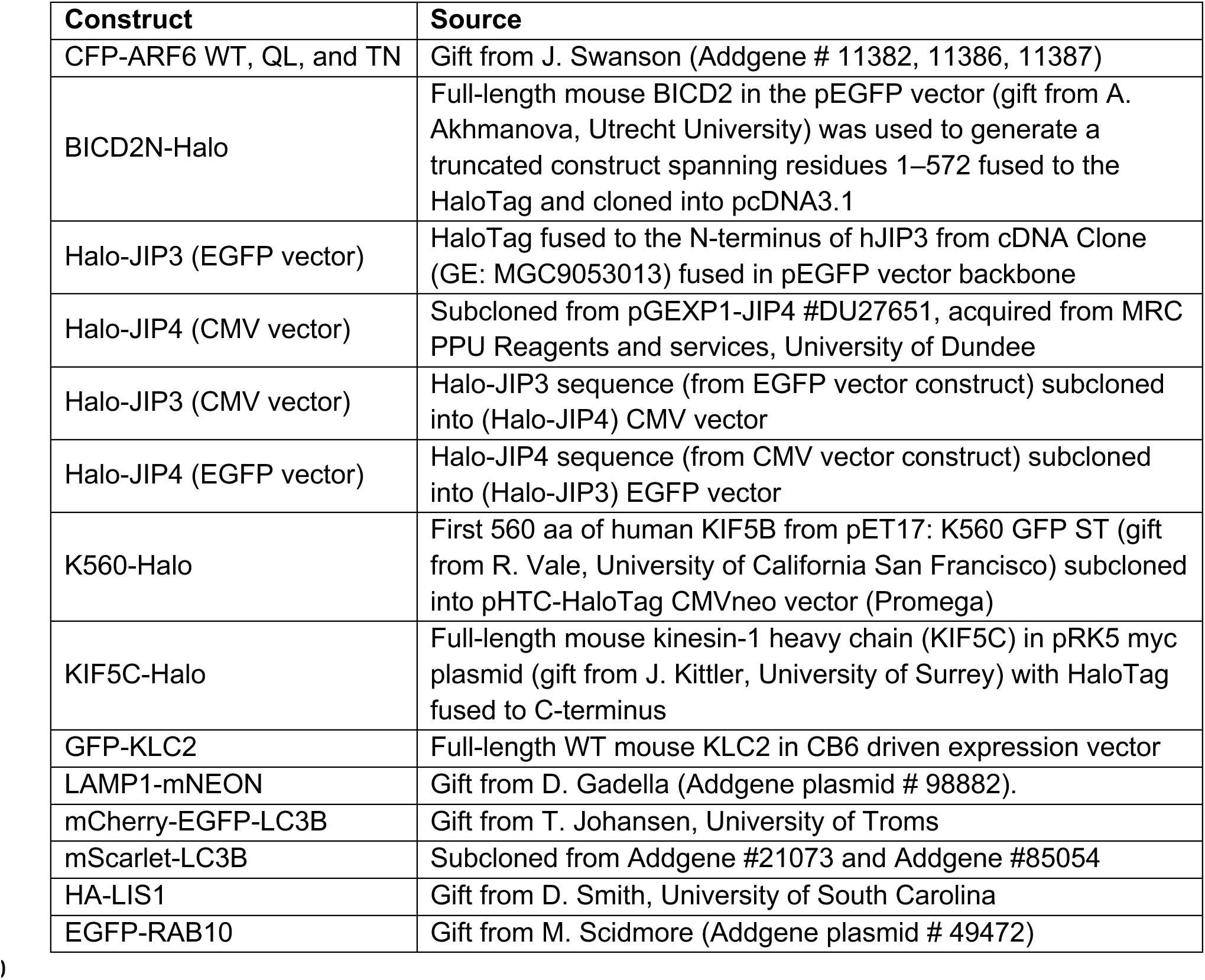

We previously used a HT-JIP4 construct in a CMVNeo backbone that expressed in cells at extremely high levels (Fig. S1, J-K). We subcloned the HT-JIP4 into an EGFP backbone (the EGFP was previously removed via subcloning), which we were already using for our HT-JIP3. This new HT-JIP4 expresses at more modest levels and does not affect AV or LAMP1 motility (Fig. S1, A-E). Therefore, we report that our previous finding was an overexpression artifact. In all neuronal experiments, we used the EGFP backbone JIP3/4; in the TIRF assays, we used the CMV backbone JIP3/4.

Antibodies include the following:

| Antibody | Source | Dilution |
| --- | --- | --- |
| anti- ACAP2/CENTB2 | ThermoFisher, PA5-18209 | WB 1:1000 |
| anti- ARF6 | Cell Signaling, D12G6 | WB 1:800 |
| anti- CYTH (pan) | ThermoFisher, MA1-062 | WB 1:1000 |
| anti- DIC | EMD Millipore, MAB1618 | PLA@1:200 |
| anti- GIT1 | Biorbyt, orb99082 | WB 1:1000 |
| anti- Halo (polyclonal) | Promega, G9281 | WB 1:500 |
| anti- Halo (monoclonal) | Promega, G9211 | WB 1:500 |
| anti- IQSEC1/BRAG2 | ThermoFisher, PA5-38019 | WB 1:500 |
| anti- IQSEC2/BRAG1 | ThermoFisher, PA5-72831 | WB 1:500 |
| anti- IQSEC3/BRAG3 | Biorbyt, orb317640 | WB 1:500 |
| anti- JIP3/MAPK8IP3 | Abcam, ab196761 | PLA 1:25 |
| anti- JIP4/SPAG9 | Cell Signaling, 5519 | PLA 1:25 |
| anti- LAMP1 | R&D Systems, AF4800 | IF 1:50 |
| anti- RAB10 | Abcam, ab237703 | WB 1:500 |
| anti- SMAP2 | Biorbyt, orb1142130 | WB 1:2000 |
| anti-Sheep (2°) 405 | Abcam, ab175676 | IF 1:1000 |
| anti-Sheep (2°) 647 | Abcam, ab150179 | IF 1:1000 |

### Primary hippocampal culture

Sprague Dawley rat hippocampal neurons at embryonic day 18 were obtained from the Neurons R Us Culture Service Center at the University of Pennsylvania. Cells (proximity ligation assay, 40,000 cells on 7mm glass; live imaging, 200,000 cells on 20 mm glass) were plated in glass-bottom 35 mm dishes (MatTek) that were precoated with 0.5 mg/ml poly-L-lysine (Sigma Aldrich). Cells were initially plated in Attachment Media (MEM supplemented with 10% horse serum, 33 mM D-glucose, and 1 mM sodium pyruvate) which was replaced with Maintenance Media (Neurobasal [Gibco] supplemented with 33 mM D-glucose, 2 mM GlutaMAX (Invitrogen), 100 units/ml penicillin, 100 mg/ml streptomycin, and 2% B-27 [ThermoFisher]) after 5-20 h. Neurons were maintained at 37 C in a 5% CO2 incubator; cytosine arabinoside (Ara-C; final conc. 1 µM) was added the day after plating to prevent glia cell proliferation. For transfections, neurons (7-10 DIV) were transfected with 0.35–1.5 µg of total plasmid DNA using Lipofectamine 2000 Transfection Reagent (ThermoFisher, 11668030) and incubated for 18-48 h.

### Live neuron imaging and analysis

Neurons were imaged in Imaging Media (HibernateE [Brain Bits] supplemented with 2% B27 and 33 mM D-glucose). Autophagosome (1-1.3 frames/sec) and lysosome (0.5-0.65 frames/sec) behavior was monitored in the proximal axon (<200 µm from the soma) of 8-12 DIV neurons for 2-3 min. Neurons were imaged in an environmental chamber at 37°C on a Perkin Elmer UltraView Vox spinning disk confocal on a Nikon Eclipse Ti Microscope with an Apochromat 100 x 1.49 numerical aperture (NA) oil-immersion objective and a Hamamatsu EMCCD C9100-50 camera driven by Volocity (PerkinElmer). Only cells expressing moderate levels of fluorescent proteins were imaged to avoid overexpression artifacts or aggregation. It should be noted that the quality of the primary neuron dissections affected autophagosomal motility, but compared conditions were always collectected from the same dissections and imaging sessions.

Kymographs were generated in ImageJ (https://imagej.net/ImageJ2) using the MultiKymograph plugin (line width, 1-5) and analyzed in ImageJ. Puncta were classified as either anterograde (moving ≥10µm towards the axon tip), retrograde (moving ≥10µm towards the soma), or stationary/bidirectional (net movement <10µm during the video). Because fluorescent LC3 is cytosolic (as well as punctate) and neurites occasionally crossed in culture, raw videos were referenced throughout kymograph analysis to ensure only real puncta (≥ 1.5 SD from the axon mean) were included in analyses. All comigration analyses were performed using kymographs.

### Proximity ligation assay

Neurons were transfected (Lipofectamine 2000) with 0.3 µg EGFP plasmid (for GFP fill) and 0.5 µg Halo-tagged effector following above protocol then 24 h later (DIV 7–8) fixed in PBS containing 4% paraformaldehyde and 4% sucrose for 8 min. Duolink^TM^ In Situ PLA Mouse/Rabbit kit with red detection reagents (Sigma-Aldrich, DUO92101-1KT) was used according to manufacturer’s protocol. We used dynein intermediate chain antibody (Mouse MAB1618) plus JIP3 antibody (Rabbit ab196761), JIP4 antibody (Rabbit Cell Signalling, 5519), or no second 1° antibody (negative control). Both 2° antibodies (Mouse and Rabbit) were added for all experiments (including negative control). Z-stacks (0.25 µm steps) were acquired on an inverted epifluorescence microscope (DMI6000B; Leica) with an Apochromat 63 x 1.4 NA oil-immersion objective and a charge-coupled device camera (ORCA-R2; Hamamatsu Photonics) using LAS-AF software (Leica). Puncta were counted manually using ImageJ.

### Cell line culture

COS-7 (ATCC) cells were maintained in DMEM (Corning) supplemented with 1% GlutaMAX and 10% FBS. Cells were maintained at 37 C in a 5% CO2 incubator. For motility assays and co-immunoprecipitation experiments, COS-7 cells were plated on 10 cm plates and transfected 24h prior to lysis using FuGENE 6 (Promega; 6-12 µg total DNA). Cells were routinely tested for mycoplasma using a MycoAlert detection kit (Lonza, LT07). COS-7 cells were authenticated by ATCC.

### Motility assay

The movement of JIP3-, JIP4-, BICD2N-, or K560-containing complexes from cell extracts was tracked using TIRF microscopy. Motility assays were performed in flow chambers constructed with a glass slide and a coverslip silanized with PlusOne Repel-Silane ES (GE Healthcare), held together with vacuum grease to form a ∼10 μl chamber. Rigor kinesin-1E236A (0.5 µM) was non-specifically absorbed to the coverslip 73 and the chamber was then blocked with 5% pluronic F-127 (Sigma-Aldrich). 250 nM GMPCPP microtubule (MT) seeds, labeled at a 1:40 ratio with HiLyte Fluor 488 tubulin (Cytoskeleton, Denver, CO), were flowed into the chamber and immobilized by interaction with rigor kinesin-1E236A.

11.25 µM free tubulin (labeled at a 1:20 ratio with HiLyte Fluor 488 tubulin) was added with the lysate to grow dynamic microtubules from the seeds. COS-7 cells grown in 10 cm plates to 70–80% confluence expressing full-length Halo-tagged HAP1, BICD2N or HaloTag alone were labeled with TMR 18–24 h post-transfection then lysed in 100 μl lysis buffer [40 mM Hepes (pH 7.4), 120 mM NaCl, 1 mM EDTA, 1 mM ATP, 0.1% Triton X-100, 1 mM PMSF, 0.01 mg ml-1 TAME, 0.01 mg ml-1 leupeptin, and 1 μg ml-1 pepstatin-A]. Cell lysates were clarified by centrifugation (17,000g) and diluted in P12 motility buffer [12 mM Pipes (pH 6.8), 1 mM EGTA, and 2 mM MgCl2] supplemented with 1 mM Mg-ATP, 1 mM GTP, 0.08 mg ml-1 casein, 0.08 mg ml-1 bovine serum albumin, 2.55 mM DTT, 0.05% methylcellulose, and an oxygen scavenging system (0.5 mg ml-1 glucose oxidase, 470 U ml-1 catalase, and 3.8 mg ml-1 glucose).

All the videos (2 min, 4-5 frames s-1) were acquired at 37°C using a Nikon TIRF microscopy system (Perkin Elmer, Waltham, MA) on an inverted Ti microscope equipped with a 100× objective and an ImageEM C9100-13 camera (Hamamatsu Photonics, Hamamatsu, Japan) with a pixel size of 0.158 µm and controlled with the program Volocity (Improvision, Coventry, England).

### Motility assay analysis

At least 5 microtubules per video were analyzed by generating kymographs using the MultiKymograph plugin of ImageJ and analyzed in Excel (Microsoft, Redmond, WA). MY polarity was determined one of two ways. (1) MT were imaged at 10 sec intervals during the entire acquisition (2 min). (2) MT were imaged for 30 sec at 4-5 frames sec -1 before and after motor imaging. In this case, only MT present in both the before and after videos were analyzed. The MT length (to which the number of events was normalized) was either (1) the final length at the end of the entire (2 min) acquisition; or (2) the initial length at the beginning of the “after” video, respectively. In either case, only non-bundled MT that could be clearly seen both growing and catastrophing regularly were analyzed.

At least 5 microtubules were analyzed per replicate; 3 biological and technical replicates were performed for a final n = 20 microtubules per condition. Kymographs were generated using the MultiKymograph plugin (line width, 1) in ImageJ (https://imagej.net/ImageJ2). Analysis was performed using KymoButler 74 with manual post-hoc curation, as described here. To be classified as an event, the duration must be greater than 0.8 seconds or the run length greater than 1.6µm, and at least 1.5 SD above the local background (surrounding ∼100µm2). To be classified as plus-end- or minus-end-directed run, the punctum must travel greater than 5 pixels (0.8µm) in that direction.

### Autophagosome fractionation

Enriched autophagosome fractions were isolated from mouse brain via sequential ultracentrifugation, adding Gly-Phe-β-naphthylamide to inactivate and deplete lysosomal vesicles and thus enhance the integrity of autophagosome-associated proteins 75; detailed protocols and validations can be found in 60. Briefly, brains were collected from wildtype mice on the C57BL/6J background (Ref 14699058) and homogenized in a tissue grinder in an ice cold buffered 10mM Hepes, 1mM EDTA, 250 mM sucrose solution, then subjected to three differential centrifugations through Nycodenz and Percoll discontinuous gradients to isolate vesicles of the appropriate size and density. The autophagosome enriched fraction was then divided and either immediately lysed for the identification of all internal and externally-associated proteins on autophagosomes (A fraction), treated with 10 μg proteinase-K for 45min at 37°C to degrade externally associated proteins and enrich for membrane-protected autophagosome cargo (P fraction), or membrane permeabilized by the addition of 0.2% triton x-100 prior to proteinase K treatment to confirm proteinase K efficacy (T fraction). The lysis buffer used contained a final concentration of 0.5% NP-40 with 1x protease and phosphatase inhibitors, PMSF and Pepstatin A. Protein concentration was measured by Bradford assay and equal amounts of protein in denaturing buffer were run on SDS-PAGE gels.

### Immunoblotting

For fluorescence Western blotting, samples were analyzed by SDS-PAGE and transferred onto PDVF Immobilon FL (Millipore). Membranes were dried for at least 1 h, rehydrated in methanol, and stained for total protein (LI-COR REVERT Total Protein Stain). Following imaging of the total protein, membranes were destained, blocked for 5min in EveryBlot Blocking Buffer (BioRad #12010021), and incubated overnight at 4°C with primary antibodies diluted in EveryBlot Blocking Buffer. Membranes were washed four times for 5 min in 1xTBS Washing Solution (50 mM Tris-HCl pH 7.4, 274 mM NaCl, 9 mM KCl, 0.1% Tween-20), incubated in secondary antibodies diluted in EveryBlot Blocking Buffer with 0.01% SDS for 1 hr, and again washed four times for 5 min in the washing solution. Membranes were immediately imaged using an Odyssey CLx Infrared Imaging System (LI-COR). Band intensity was measured in the LI-COR Image Studio application.

### Analysis of organelle enrichment publications

Goldsmith et al., (2022) and Dumrongprechachan et al., (2022) performed AV and lysosomal enrichments, respectively, and performed mass spectrometry on the resultant proteins to determine proteins associated with the organelles. To compare these datasets, which pulled down different amounts of total protein, we normalized the number of peptides from each to the average number of peptides detected for ARF-related proteins (normalization factor) in each preparation. The number of peptides for each detected protein was divided by the normalization factor (13.5 for lysosomes, 2.6 for AVs). Note that for the AVs, we used the number of peptides in the fraction not treated with proteinase K (AV fraction).

### Statistics

All statistical analyses were performed in Prism (GraphPad, San Diego, CA). Bars represent mean ± unless otherwise indicated. n indicates the number of events or cells pooled across at least 3 trials per experiment. Parametric or nonparametric tests were used where appropriate, but formal testing was not performed. Statistical measures are described in the legends.

## Author contributions

Sydney E. Cason, Conceptualization, Resources, Data curation, Formal analysis, Validation, Investigation, Visualization, Methodology, Project administration, Writing—original draft and review/editing; Erika L.F. Holzbaur, Conceptualization, Supervision, Funding acquisition, Project administration, Writing—review/editing

## Supplement legends

**Figure S1. JIP3 or JIP4 overexpression does not affect AV or lysosome transport. (A-C)** Example kymographs showing LC3 and LAMP1 puncta motile behavior in the axons of neurons expressing HaloTag (HT) alone (Tag), HT-JIP3, or HT-JIP4. Annotated kymographs (annot.) show paths pseudo-colored for visualization; heavier weight lines represent paths with both LC3 and LAMP1 co-migrating. **(D-E)** Quantification of the fraction of LC3 or LAMP1 puncta moving retrograde (≥10µm towards the soma), anterograde (≥10µm towards the axon tip), or exhibiting bidirectional/stationary motility (moving <10µm). Symbols indicate comparison to Tag; n = 15 neurons; two-way ANOVA with Tukey’s multiple comparisons test; LC3 [variation between motile fractions (P < 0.0001) but no variation between conditions (P > 0.9999) nor interaction (P = 0.4514)]; LAMP1 [variation between motile fractions (P < 0.0001) and mild interaction (P = 0.0032) but no variation between conditions (P > 0.9999)]. **(F-G)** LC3 and LAMP1 puncta density (per µm in a 2 min video) in cells expression HT-JIP3, HT-JIP4, or Tag. n = 11-15 neurons; one-way ANOVA with Tukey’s multiple comparisons test (LC3, P = 0.9076; LAMP1, P = 0.9397). **(H-I)** Colocalization between LC3 and LAMP1 puncta in cells expression HT-JIP3, HT-JIP4, or Tag. n = 11-15 neurons; one-way ANOVA with Tukey’s multiple comparisons test (LC3, P = 0.3154; LAMP1, P = 0.5569). **(J-K)** Western blot and quantification demonstrating that HT-JIP3 or JIP4 in the CMV backbone expresses far more highly than in the EGFP backbone (EGFP sequence has been removed by subcloning). This experiment was performed using COS-7 cells, which we transfected at the same confluence (∼50%) with FuGene and equal DNA quantities. After lysis in RIPA buffer, we assessed for protein concentration using BCA assay. Equal protein concentrations were loaded, which was confirmed using Revert Total Protein Stain. Finally, a monoclonal HT antibody was used to assess expression of the HT proteins. n = 3; one-way ANOVA with Sidak’s multiple comparisons test (JIP3 EGFP v. JIP3 CMV, P = 0.0051; JIP4 EGFP v. JIP4 CMV, P = 0.0444; JIP3 EGFP v. JIP4 EGFP, P = 0.9118; JIP3 CMV v. JIP4 CMV, P = 0.0857). **(L)** Example PLA negative (Neg.) control (missing JIP3/4 antibody). **(M)** Quantification of DIC PLA puncta either with JIP3, JIP4, or no second 1° antibody (Neg. control). n = 20-21 neurons; one-way ANOVA with Tukey’s multiple comparisons test (JIP3 v. JIP4, P = 0.6763; JIP3 v. Neg., P < 0.0001; JIP4 v. Neg., P < 0.0001).

**Figure S2. JIP3 and JIP4 do not induce kinesin activity *in vitro*. (A)** Example kymograph of labeled KIF5C^1-560^ (K560) activity. **(B)** Number of events (per µm microtubule per min) observed for K560-containing complexes. n = 20 MT each; Kruskal-Wallis test with Dunn’s multiple comparisons; K560: 0 v. –, P = 0.0004; 0 v. +, P = 0.3092; – v. +, P < 0.0001. **(C-E)** Quantification of velocities and run lengths towards the MT plus end for K560-, JIP3-, or JIP4-containing complexes. All velocity histograms were fit to a Gaussian curve and all run length histograms (1– cumulative distribution frequency) were fit to a one phase decay. Listed values are median (25^th^ percentile-75^th^ percentile). JIP3/JIP4, n = 19-22 events; K560, n = 160 events. **(F-G)** Quantification of the directionality of runs on each microtubule. Runs were defined as events ≥ 0.8 µm in length towards either the minus-or plus-end of the microtubule. Note that the + LIS1 and – KIF5C & KLC2 are repeated from Figure 2. n = 15-20 MT each; Kruskal-Wallis test with Dunn’s multiple comparisons; LIS1: JIP3 P > 0.9999, JIP4 P > 0.9999; KIF5&KLC: JIP3 P = 0.3572, JIP4 P > 0.9999.

**Figure S3. RAB10 expression does not affect the formation of JIP3/4-dynein complexes. (A-B)** Quantification of mSc-LC3 puncta linear density and colocalization with LAMP1-HT. n = 9 neurons; unpaired t test; density, P = 0.7068; colocalization, P = 0.3086. **(C-D)** Quantification of LAMP1-HT puncta linear density and colocalization with mSc-LC3. n = 9 neurons; unpaired t test; density, P = 0.0438; colocalization, P = 0.8564. **(E-F)** Total JIP3-or JIP4-DIC PLA puncta linear density, compared between cells expressing EGFP alone (Tag) and cells expressing EGFP-RAB10. Note that the Tag data is repeated from Fig. S1 M. n = 20 neurons; unpaired t test; JIP3, P = 0.3278; JIP4, P = 0.5541. **(G-H)** Colocalization between JIP3/4-DIC PLA puncta and AVs or lysosomes. Note that Note that the Tag data is repeated from Fig. 1 K, M. n = 20 neurons; one-way ANOVA with Tukey’s multiple comparisons test; JIP3 (LC3 only, P = 0.7989; LC3 + LAMP1, P = 0.9971; LAMP1 only, P = 0.7486); JIP4 (LC3 only, P = 0.9984; LC3 + LAMP1, P = 0.2160; LAMP1 only, P > 0.9999).

**Figure S4. ARF6 expression does not affect LC3 or LAMP1 density or colocalization. (A)** Quantification of mCh-LC3 puncta linear density. n = 13-16 neurons; one-way ANOVA with Sidak’s multiple comparisons test; WT v. QL, P = 0.6382; WT v. TN, P = 0.8339; QL v. TN, P = 0.6382. **(B)** Quantification of mCh-LC3 colocalization with LAMP1-HT. one-way ANOVA with Sidak’s multiple comparisons test; n = 9-11 neurons; WT v. QL, P = 0.4183; WT v. TN, P = 0.9791; QL v. TN, P = 0.4183. **(C)** Quantification of LC3 non-processive movement, as described by 𝚫 run length (net run length of each vesicle subtracted from its total run displacement). n = 71-111 puncta; Kruskal-Wallis test with Dunn’s multiple comparisons test; WT v. QL, P > 0.9999; WT v. TN, P > 0.9999; QL v. TN, P > 0.9999. **(D)** Quantification of LAMP1-HT puncta linear density. one-way ANOVA with Sidak’s multiple comparisons test; n = 10 neurons; WT v. QL, P = 0.1665; WT v. TN, P = 0.6735; QL v. TN, P = 0.2495. **(E)** Quantification of LAMP1-HT colocalization with mCh-LC3. one-way ANOVA with Sidak’s multiple comparisons test; n = 10 neurons; WT v. QL, P = 0.6020; WT v. TN, P = 0.5742; QL v. TN, P = 0.3846. **(F)** Quantification of LAMP1 non-processive movement, as described by 𝚫 run length. n = 13-16 neurons; Kruskal-Wallis test with Dunn’s multiple comparisons test; WT v. QL, P > 0.9999; WT v. TN, P > 0.9999; QL v. TN, P > 0.9999.

**Figure S5. ARF6 does not affect motile events *in vitro*. (A)** Example kymographs showing the activity of JIP3-and JIP4-containing complexes in the presence of CFP-ARF6^T27N^. **(B-C)** Quantification of the number of total landing events for JIP3- and JIP4-containing complexes in the presence of CFP-Arf6^Q67L^ or CFP-Arf6^T27N^. n = 20-23 MT each; unpaired t test; JIP3, P < 0.6856; JIP4, P = 0.1028. **(D-E)** Number of events (per µm microtubule per min) observed for JIP3-containing and JIP4-containing complexes in the presence of CFP-Arf6^Q67L^ or CFP-Arf6^T27N^. Complexes with a net direction of “0” were stationary landing events, while complexes with a net direction of “–” or “+” moved ≥ 0.8 µm towards the minus- or plus-end of the microtubule respectively. Note that the Arf6^QL^ data from is repeated from Figure 8. Kruskal-Wallis test with Dunn’s multiple comparisons. n = 20-23 MT each. JIP3 QL v. TN: 0, > 0.9999; –, P > 0.9999; +, P > 0.9999. JIP4 QL v. TN: 0, 0, > 0.9999; –, P = 0.9389; +, P > 0.9999. Dashed lines indicate mean without added ARF6. **(F)** Quantification of the activity of JIP3 or JIP4-containing dynein complexes in the presence of CFP-Arf6^Q67L^. **(G)** Quantification of the activity of JIP3 or JIP4-containing kinesin complexes in the presence of CFP-Arf6^Q67L^. All velocity histograms were fit to a Gaussian curve and all run length histograms (1– cumulative distribution frequency) were fit to a one phase decay. Listed values are median (25^th^ percentile-75^th^ percentile). n = 39-140 events.

