## Supplementary figures and images for "Axonal transport of autophagosomes is regulated by dynein activators JIP3/JIP4 and ARF/RAB GTPases"

### Supplemental Figures

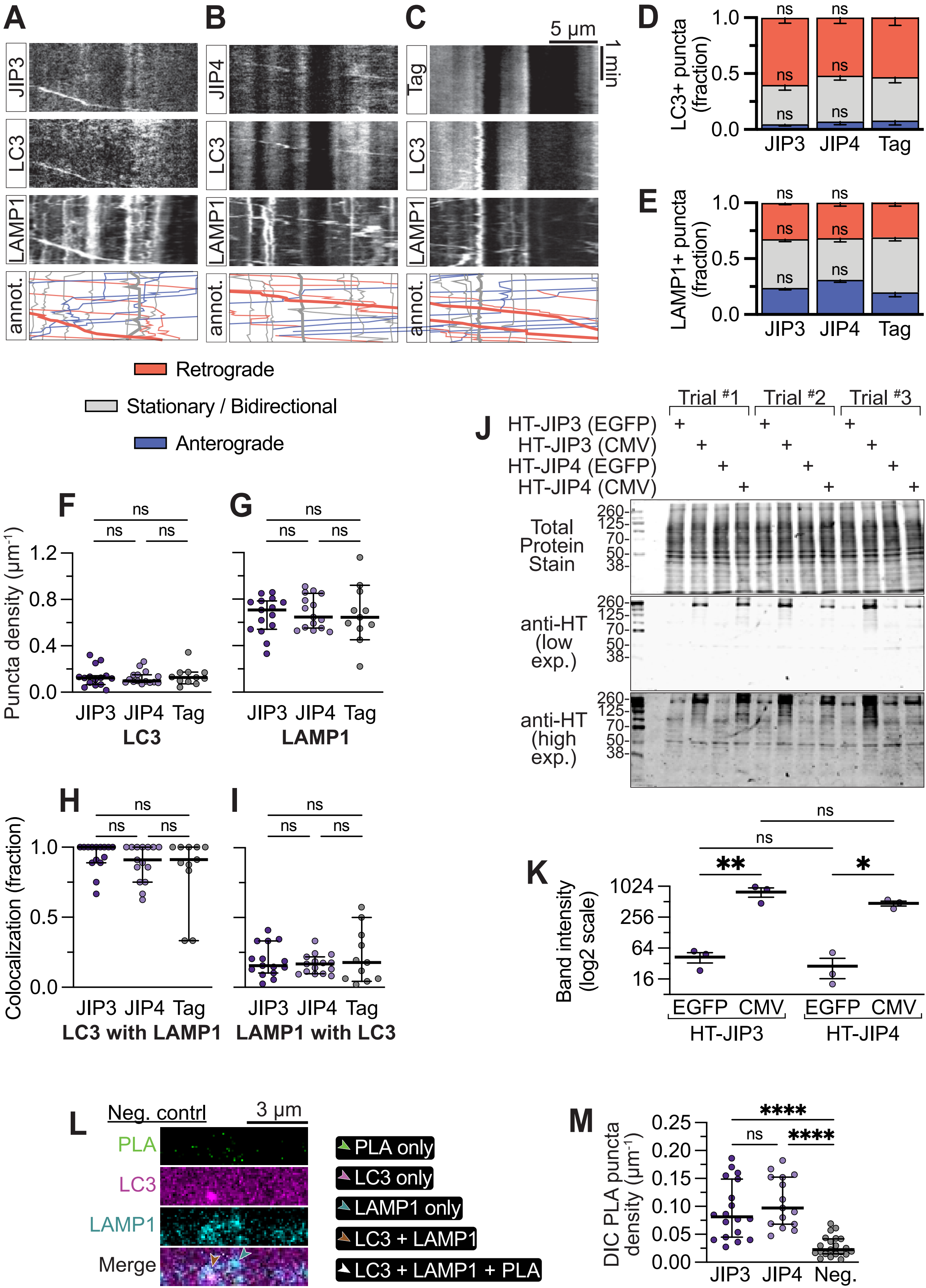

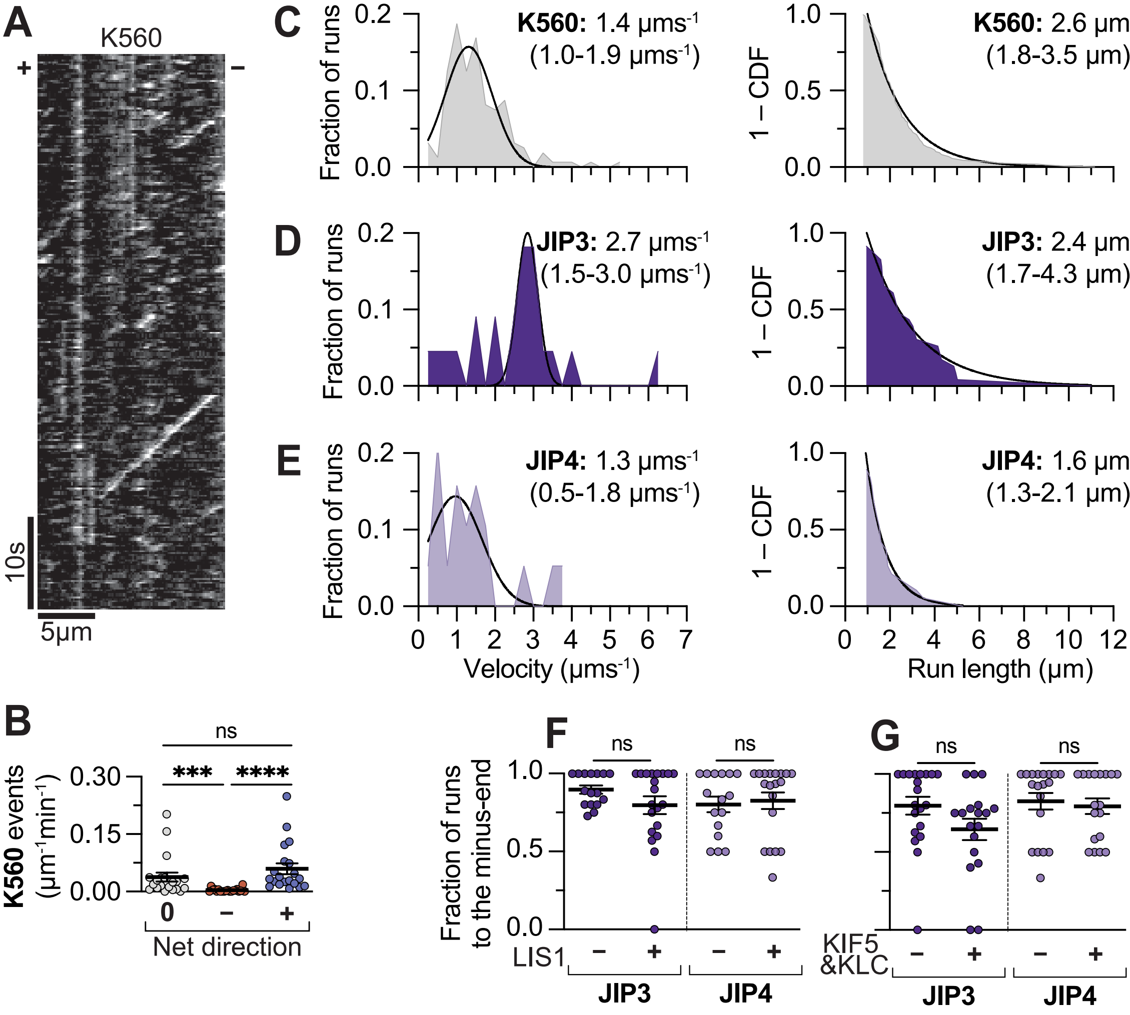

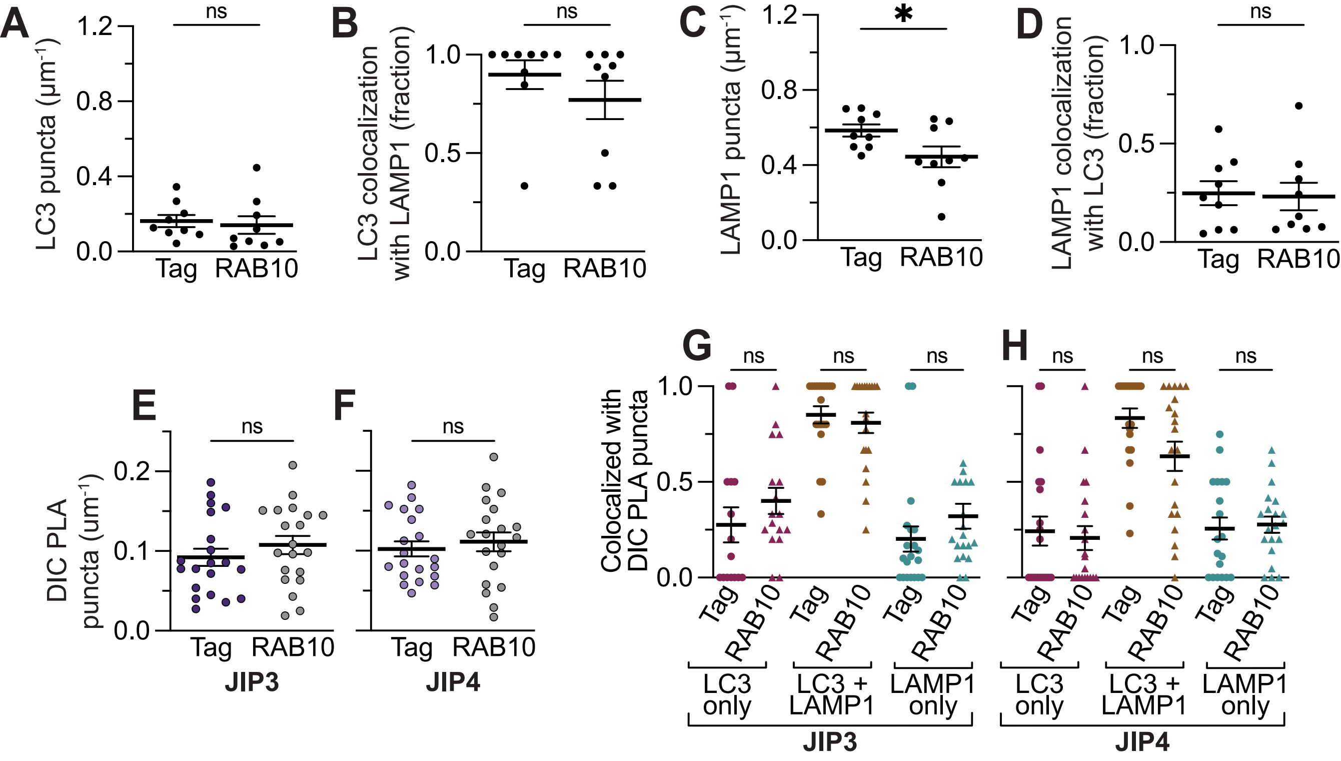

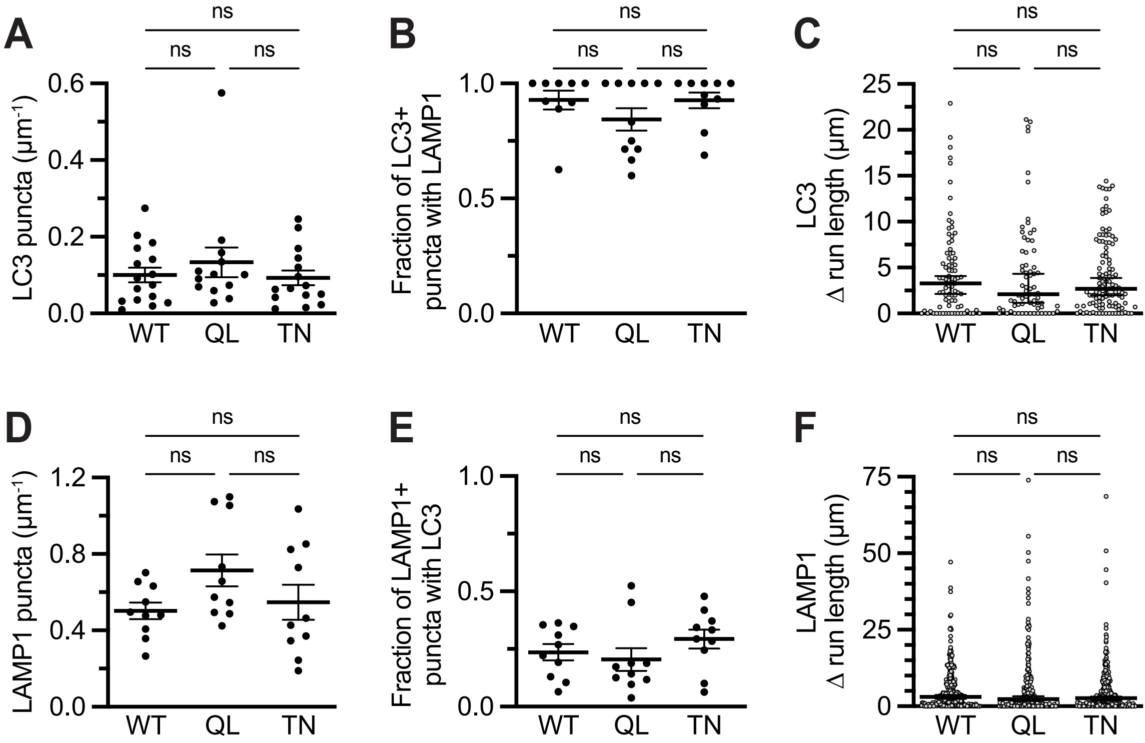

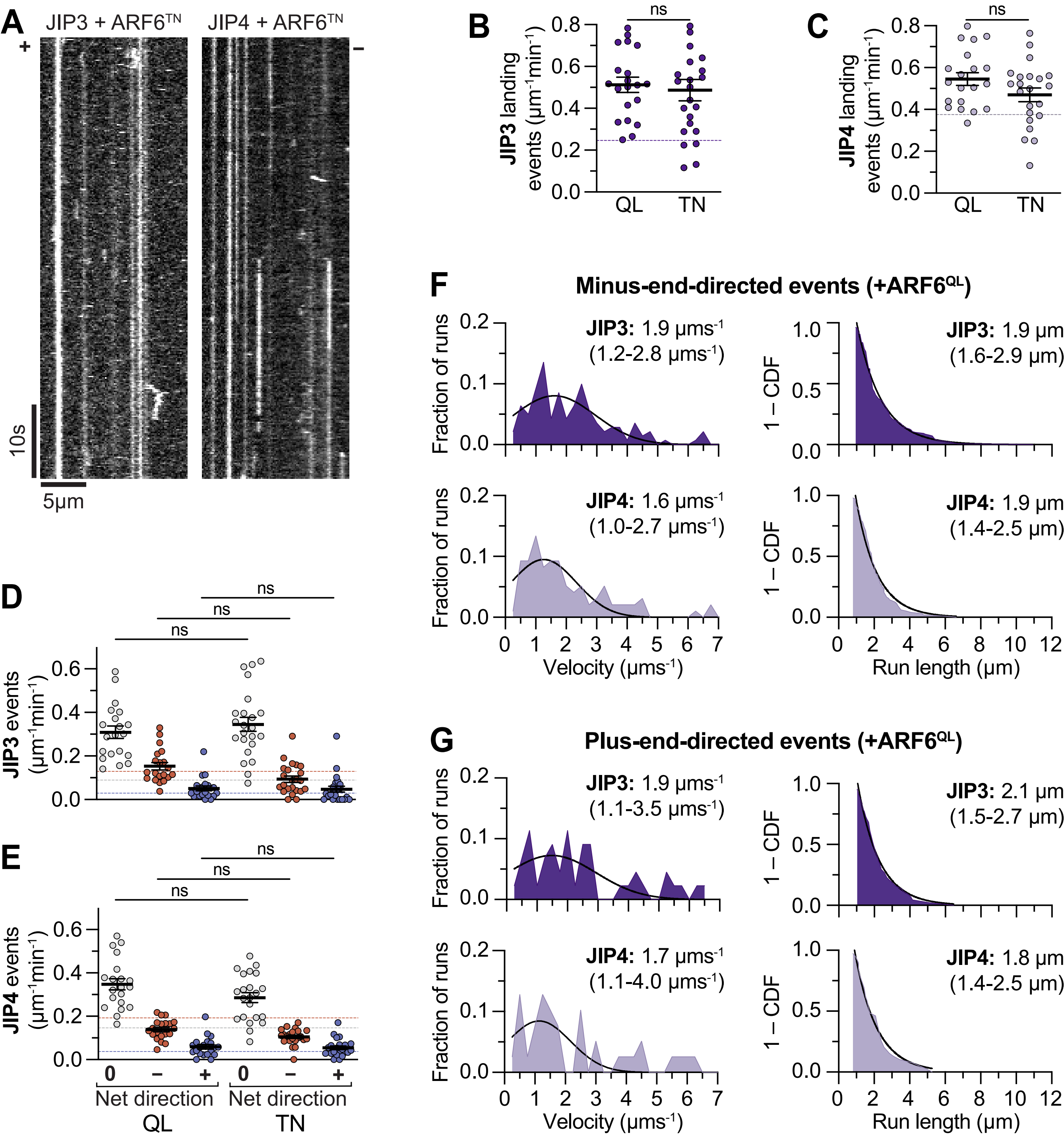
